# Quantifying the interplay between rapid bacterial evolution within the mouse intestine and transmission between hosts

**DOI:** 10.1101/2020.12.04.412072

**Authors:** Kimberly S. Vasquez, Lisa Willis, Nate Cira, Katharine M. Ng, Miguel F. Pedro, Andrés Aranda-Díaz, Manohary Ranjendram, Feiqiao Brian Yu, Steven K. Higginbottom, Norma Neff, Gavin Sherlock, Karina B. Xavier, Stephen Quake, Justin Sonnenburg, Benjamin H. Good, Kerwyn Casey Huang

## Abstract

Due to limitations on high-resolution strain tracking, selection dynamics during gut-microbiota colonization and transmission between hosts remain mostly mysterious. Here, we introduced hundreds of barcoded *Escherichia coli* strains into germ-free mice and quantified strain-level dynamics and metagenomic changes. Mutants involved in motility and utilization of abundant metabolites were reproducibly selected within days. Even with rapid selection, coprophagy enforced similar barcode distributions across co-housed mice. Whole-genome sequencing of hundreds of isolates quantified evolutionary dynamics and revealed linked alleles. A population-genetics model predicted substantial fitness advantages for certain mutants and that migration accounted for ~10% of the resident microbiota each day. Treatment with ciprofloxacin demonstrated the interplay between selection and transmission. While initial colonization was mostly uniform, in two mice a bottleneck reduced diversity and selected for ciprofloxacin resistance in the absence of drug. These findings highlight the interplay between environmental transmission and rapid, deterministic selection during evolution of the intestinal microbiota.

## Introduction

The gut microbiota is important for many aspects of host physiology, including resistance to pathogen invasion (Litvak and Baumler, 2019), modulation of the immune system (Pickard et al., 2017), and metabolism (Visconti et al., 2019). The close relationship between mammalian hosts and microbes has emerged through extensive co-evolution (Moeller et al., 2016). While substantial progress has been made in correlating microbiota composition to host health and disease, the complexity and rapid turnover of this ecosystem has limited our understanding of the dynamics of colonization, as well as of the magnitude and importance of transmission between hosts and seeding from the environment.

Comprehensive studies of genetic variability within commensal species have revealed substantial strain-level variation (Poyet et al., 2019) that potentially reflects a range of adaptations. Diet has been associated with particular strains of the commensal *Prevotella copri* (De Filippis et al., 2019), due to mutations in polysaccharide-utilization loci (Fehlner-Peach et al., 2019). Genetic variation of the human intracellular pathogen *Listeria monocytogenes* has been connected to differences in ecology and virulence potential, leading to diagnostic applications (Ragon et al., 2008). Genetic bottlenecking of *Yersinia pestis* arises because this species can infect with only a few cells (Perry and Fetherston, 1997), illustrating the potential importance of colonization bottlenecks in genetic variability and selection. In humans, resident populations of gut bacteria exhibit a relatively clonal structure, with a few strains of each species at intermediate or high frequencies (Garud et al., 2019; Truong et al., 2017); it is currently unclear whether this phenomenon is typically caused by strong colonization bottlenecks, the transient invasion of external strains, or stable ecological sub-structure within species. Thus, a deeper understanding of the extent of selection and migration, and the time scales over which genetic variation is established within a host would have broad relevance regarding population dynamics within the gut microbiota.

Laboratory long-term evolution experiments have provided quantitative insights into the interplay between genetic variation and selection during microbial evolution. During repeated *in vitro* passaging for >60,000 generations, *Escherichia coli* experienced large increases in fitness (Wiser et al., 2013), cell size (Philippe et al., 2009), and metabolic potential (Grosskopf et al., 2016). Metagenomic sequencing revealed that microecologies (multiple co-existing strains) can persist within these evolved populations for >10,000 generations, while their relative frequencies are continually perturbed by the accumulation of additional mutations (Good et al., 2017). During *in vitro* passaging of budding yeast, the use of DNA barcodes (the genomic integration of unique sequences for subsequent identification) and sequencing to monitor the relative frequencies of many competing lineages revealed rapid and reproducible evolutionary dynamics (Levy et al., 2015), and large-scale isolation of adapted strains permitted genotype-fitness mapping (Venkataram et al., 2016) and uncovered growth-phase tradeoffs (Li et al., 2019).

Several previous studies have capitalized on a barcoding approach to study selection in the mouse gut. Colonization of conventional mice with two *E. coli* strains harboring different fluorescent proteins found ubiquitous clonal interference (competition among lineages arising from mutations that appeared independently) (Barroso-Batista et al., 2014). Evolution of *E. coli* in ex-germ-free mice was found to depend on selection based on amino-acid consumption and niche availability (Barroso-Batista et al., 2020). Clonal interference was also observed in conventional mice infected with eight *Salmonella enterica* serovar Typhimurium strains, each of which harbored a distinct DNA barcode (and were otherwise isogenic); only one or a few strains persisted in the gut after 35 days (Lam and Monack, 2014). Notably, distinct sets of barcodes emerged in individual mice, indicating either stochastic colonization, selection, or post-colonization drift (Lam and Monack, 2014). Due to the presence of other microbiota members, it is difficult to determine whether lack of barcode diversity is inherent to *Salmonella* colonization or was due to the rest of the community being present. Moreover, the individual housing of mice and/or the limited number of barcodes in these previous studies precluded measurements of the importance of transmission between hosts sharing the same environment. Perturbations such as antibiotic treatment (Ng et al., 2019) or osmotic diarrhea (Tropini et al., 2018) can apply selective pressures that lead to long-term changes in microbiota diversity and composition as well as the emergence of resistant strains (Ng et al., 2019). It has yet to be determined how such perturbations affect intraspecies genetic diversity, migration between hosts, and environmental reservoirs, which could in turn affect recovery of the microbiota and the host. Barcodes provide a powerful tool to quantify these fundamental behaviors.

Here, we utilized heritable DNA barcodes to simultaneously track, at high temporal resolution, the evolutionary dynamics and genetic adaptations of nearly 200 isogenic strains of *E. coli* in living animal hosts. Colonization was initially even across most of these mono-colonized mice, followed by multiple waves of selection. In all cages, constant migration between hosts enforced similar abundances. Strain dominance was dependent on the fitness conferred by newly acquired mutations. Using metagenomic and whole-genome sequencing, we repeatedly identified *in vivo* selection for mutations that affect motility and metabolism, and nearly identical evolutionary trajectories were identified in a replicate experiment. Metabolomic analysis of mouse intestinal and fecal contents and phenotype profiling of isolates suggested selection for mutations that enable utilization of raffinose. Using a population-genetics model, we determined that migration from environmental reservoirs accounts for a substantial fraction of the resident microbiota, which homogenizes the barcode composition across mice in the same cage and quantitatively predicts transmission kinetics. Treatment with ciprofloxacin created a large bottleneck that allowed for transmission between cages and selected for a single resistant strain, demonstrating that strain-level variation can be altered dramatically by environmental perturbations. Unexpectedly, the resistance mutation was pre-existing due to a bottleneck in a few mice during colonization. This study demonstrates the power of genetic barcoding for uncovering rates of selection and transmission across environments.

## Results

### Stable colonization of ex-germ-free mice by a barcoded *E. coli* library

To determine the kinetics of gut colonization and community assembly at the intra-species level, we sought to investigate how subpopulations of the enteric bacterium *E. coli* would colonize, compete, and evolve within gnotobiotic mice in the absence of other species. A previous study generated a library of high-copy plasmids with ampicillin and chloramphenicol resistance cassettes that each carried a unique barcode (Figure 1A) (Cira et al., 2018). We transformed the common laboratory wild-type strain MG1655 with 191 distinct barcoded plasmids. Our choice of MG1655 was motivated by the goal of investigating rapid selection, since we surmised that the lab-adapted MG1655 strain would undergo accelerated evolution within the *in vivo* environment of the mammalian gut. All transformed strains had similar growth kinetics *in vitro* (Figure S1A). We split these 191 strains into sets of 95 (set 1) and 96 (set 2) to separately colonize mice, enabling us to study both evolution within mice and the transmission of barcoded strains from one group of mice into another.

**Figure 1:**
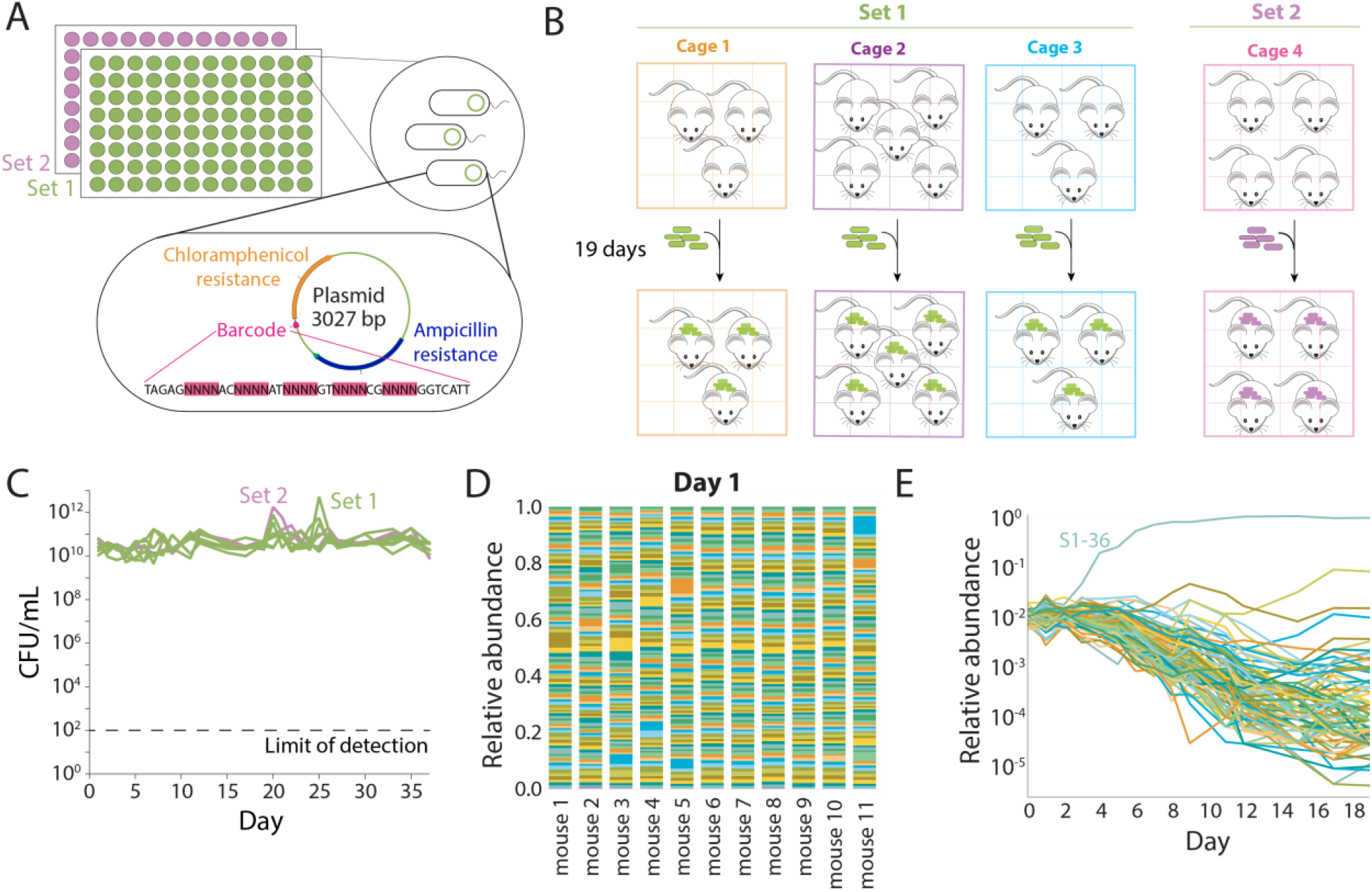
Colonization of ex-germ-free mice by a barcoded *E. coli* library is initially even, but a single strain dominates within one week. A) Plasmids containing a unique 28-bp DNA barcode and encoding resistance cassettes to chloramphenicol and ampicillin were used to transform *E. coli* MG1655, a common laboratory wild-type strain. We isolated 191 unique strains, sequenced them to verify the presence of a single, unique barcode surrounded by common primer sequences, and split them into two sets of 95 (S1) and 96 (S2) strains that were used to independently colonize ex-germ-free mice. The abundance of each strain was tracked via sequencing after barcode amplification. B) Schematic of housing layout of mice colonized with barcoded *E. coli*. Three cages with 3, 5, and 3 mice were colonized with S1 strains, and one cage with 4 mice was colonized with S2 strains. C) After the mice were colonized, fecal samples were collected daily and plated on LB with 100 μg/mL ampicillin and 35 μg/mL chloramphenicol. Mice gavaged with S1 strains were colonized to similar levels as mice gavaged with S2 strains (~10^10^-10^11^ CFUs/mL), and the bacterial load remained stable over the entire experiment. D) One day after gavage with the barcoded strains, the 11 mice gavaged with S1 strains were approximately evenly colonized with each of the 95 strains. E) A single barcode can become dominant within one week. Over the first 19 days, most strains in mouse 2 continuously decreased in relative abundance as barcode S1-36 took over after day 4, although a few maintained their level of relative abundance or experienced a resurgence after day 10.

We gavaged 11 and 4 germ-free mice on day 0 with an equal-volume mixture of 10^8^ colony forming units (CFUs) of set 1 (S1) and set 2 (S2) barcoded strains, respectively (equivalent to ~10^6^ CFUs per strain) (Figure 1B). To prevent cross-contamination, mice colonized with S1 strains were housed in separate gnotobiotic isolators (a container that maintains a germ-free environment) from mice colonized with S2 strains for the first 19 days. This time frame allowed the *E. coli* strains to fully colonize and compete for niches. Four of the S1-colonized mice were also observed for a further 18 days to identify selection on longer time scales.

Feces from each mouse were collected daily beginning on day 1 to enable barcode tracking at high temporal resolution, and DNA was extracted for barcode and metagenomic sequencing. To evaluate the necessity of antibiotic selection for plasmid maintenance, after day 7 the mouse drinking water was supplemented with 0.2 mg/mL ampicillin; ampicillin was given only after day 7, to avoid disruption of the initial colonization dynamics. We measured CFUs/mL from each fecal sample on LB plates with ampicillin and chloramphenicol and found that mice were colonized with ~10^10^-10^11^ CFUs/mL by day 1, with this level maintained thereafter (Figure 1C). There was no substantial change in CFUs/mL once ampicillin was added to the drinking water on day 7 (Figure 1C). Moreover, CFUs/mL measured on LB plates were essentially the same as on antibiotic-supplemented plates (Figure S1B). Thus, antibiotic selection is not required for retention of the plasmid *in vivo*.

Barcode amplification and sequencing enabled quantification of the relative abundances of each barcode throughout the experiment. The initial loss of some strains would represent a strong bottleneck during colonization. However, we found that every barcode was present in S1 mice in the day 1 samples, with an approximately even distribution of relative abundances in all mice (mean coefficient of variation of 0.44; Figure 1D). In a representative mouse, this even distribution was maintained for several days, before apparent selection of a more fit strain (S1-36) starting on day 4 (Figure 1E). Thus, we conclude that the entire set of barcodes transited through the mouse intestinal tract and colonized with approximately equal fitness.

### Co-housed mice maintain highly similar barcoded strain abundances despite rapid adaptation within hosts

Similar to previous studies (Barroso-Batista et al., 2014), we observed rapid expansion of individual barcodes starting on day 4, suggesting that some lineages had started to acquire adaptive mutations. Given our co-housing design, we sought to determine whether these barcodes would expand independently in different mice (e.g. due to independent mutation events), or whether the transmission of newly acquired mutations would lead to similar barcode trajectories within the same cage. In the first cage of S1 mice, the same strain (S1-36) rapidly expanded in all mice by day 4 (Figure 2A, S2A). This barcode also rapidly expanded in the other two cages of S1 mice, suggesting that its fitness benefit was potentially caused by a pre-existing mutation that was present at low frequency in the initial stock (Figure 2B, S2B). However, the other two strains that reached >1% frequency in cage 1 (S1-9 and S1-57) expanded at similar times in all mice in the cage, but did not expand at all in the other two cages (Figure 2A, S2A). This observation suggests that strain transmission between co-housed mice (rather than the highly unlikely independent emergence of the same barcodes in each mouse) was an important factor in establishing community composition *in vivo*.

**Figure 2:**
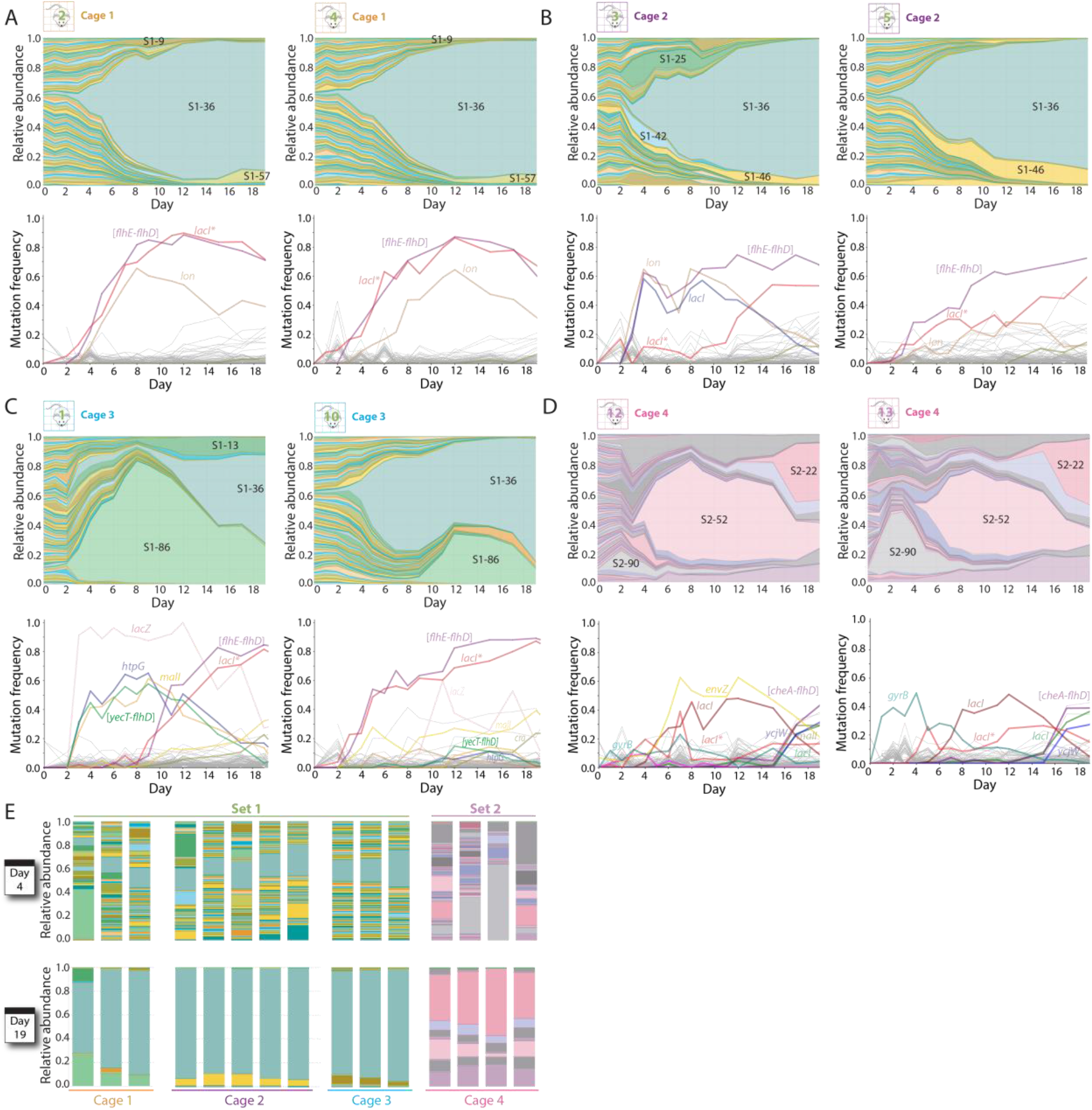
Cage-specific subsets of barcoded strains quickly expand in abundance due to selection of mutations in motility- and metabolism-related genes. A-C) Relative abundances of barcodes (top) and mutations (bottom) in two representative mice within each cage of S1 mice. In all three cages, S1-36 took over; this strain harbors mutations in *lacI* and large motility deletions such as [*flhE-flhD*]. D) Relative abundances of barcodes (top) and mutations (bottom) in two representative S2 mice. Multiple waves of partial barcode takeover occurred, with several barcodes persisting at relative abundance >5% by day 19. E) On day 4, relative abundances between mice in the same cage were much more variable compared to those on day 19, despite differences among cages.

Metagenomic sequencing allowed us to link barcode dynamics with the emergence of specific mutations (Table S1). All mice in cage 1 exhibited high frequencies of three mutations: a 4-bp insertion in *lacI*, which encodes the repressor of the Lac operon (henceforth *“lacI*”*); a 15,897-bp deletion in the flagellar operon from *flhE* to *flhD* (henceforth *“[flhE-flhD]”*), and a mutation in *lon*, which encodes the Lon protease. All three mutations reached >60%, indicating that they were present in S1-36 and were potentially drivers of the dominance of this strain (Figure 2A, S2A).

Identical *lon* and [*flhE-flhD*] alleles were also observed during the expansion of S1-36 barcode in cage 2, suggesting a common mutational origin (Figure 2B, S2B); these mutations were not detectable from metagenomic sequencing of the original S1-36 stock. However, in one of the mice from this cage (mouse 3), these alleles were initially accompanied by a different mutation in *lacI* (a point mutation, G272V) suggesting that the *lacI* mutations likely occured after the initial [*flhE-flhD*] and *lon* mutations. Interestingly, the *lacI^G272V^* mutation in mouse 3 was eventually outcompeted by *lacI\** by day 20 (Figure 2B), demonstrating within-barcode competition and highlighting the synergy between metagenomic and barcode sequencing data. In another mouse (#5), the *lacI\** allele increased as in cage 1, suggesting that *lacI\** has higher fitness than *lacI^G272V^* (Figure 2B).

In the third cage of S1 mice, again S1-36 was the most abundant strain on day 19 (Figure 2C, S2C). However, in one mouse (#1), another strain (S1-86) first transiently became dominant until day 8, followed by slow replacement by S1-36 (Figure 2C, left). In another mouse from the same cage (#10), S1-36 expanded first, but then S1-86 began to increase on day 9 (Figure 2C, right), directly after the peak in abundance in mouse 1, suggesting that the beneficial variant of S1-86 was transmitted from mouse 1 to mouse 10. Strain S1-86 harbored a *lacZ* point mutation, emphasizing the role of the lactose operon in mediating adaptation in this environment (Figure 2C, left). Mouse 1 contained high frequencies of the *lacI\** and [*flhE-flhD*] alleles associated with S1-36, as in cage 1 (Figure 2C). However, the initial expansion of S1-86 allowed us to identify the mutations associated with this strain: a 16,203-bp motility deletion ([*yecT-flhD*]) as well as mutations in *htpG* and *malI* (which encodes the repressor of the maltose-utilization operon). The data from this cage suggest a general fitness benefit of large motility deletions, as well as the importance of the combination of the *lacI*\* and [*flhE-flhD*] alleles for S1-36 to outcompete S1-86. After S1-86 invaded mouse 10, it was again outcompeted at the same time that it was outcompeted in the other two mice. coincident with the emergence of a *cra* mutation arose, presumably in S1-36 (Figure 2C, S2C).

In two of the four S2 mice (#12 and #14), the initial abundances of the 96 barcoded strains were approximately uniformly distributed (Figure S2D), as in the S1 mice. However, in the other two S2 mice (#13 and #15), the initial distribution of abundances was highly uneven, with five barcodes having >15% relative abundance (Figure S2D); in mouse 15, strains S2-5, S2-33, and S2-90 collectively accounted for >96% of the total bacteria on day 1 (Figure S2D). Nonetheless, by day 4, the evenness of barcodes abundances returned to essentially the same level as in the other two S2 mice (Figure 2E), suggesting that exchange of bacteria between mice effectively eliminated the initial unevenness. Moreover, the abundances of all strains selected for by the bottleneck quickly decreased after day 1 (Figure 2D, S2E). These dynamics suggest that in mice 13 and 15 the bacteria faced a strong but short-lived selective pressure specific to initial colonization, and mutations that conferred fitness benefits in the face of this selective pressure carried high pleiotropic fitness costs directly afterward.

In the cage of S2 mice, five barcodes reached at least 10% at some point, and seven barcodes collectively dominated the population by day 19 (Figure 2D, S2E). As in the S1 mice, we observed the emergence of metabolic and motility-related mutations, including two *lacI* mutations distinct from *lacI*\* and *lacI^G272V^* (Figure 2D, S2E). The genetic variation exhibited by S2 strains suggests a wide diversity of adaptive mutations that likely did not emerge in S1 mice due to the dominance of the *lacI*\* and [*flhE-flhD*] allele combination in S1 mice. Nonetheless, by day 19 the compositions of all S2 mice were highly similar to each other (Figure 2E), with Pearson correlation coefficients of 0.95-0.99 between the pairs of mice (Figure S2F). These data indicate the capacity of a host to support multiple strains despite multiple waves of expansion and competition in the first 19 days, with transmission helping to establish a common composition among co-housed mice. These findings emphasize the importance of environmental reservoirs in shaping community composition among co-housed mice.

### Flagellar motility and carbon utilization are reproducibly targeted in the gut

Since S1-36 developed a strong fitness advantage within a short time, we repeated the experiment, gavaging three new cages of germ-free mice (*n*=9 total) with the original S1 barcode pool except S1-36, to ensure a different evolutionary trajectory; we refer to these as S1^36-^ mice. All S1^36-^ barcodes were detected in the mice on day 1 (Figure 3A), and the library colonized to similar CFUs/mL (Figure S3A) as in our first experiment (Figure 1C).

**Figure 3:**
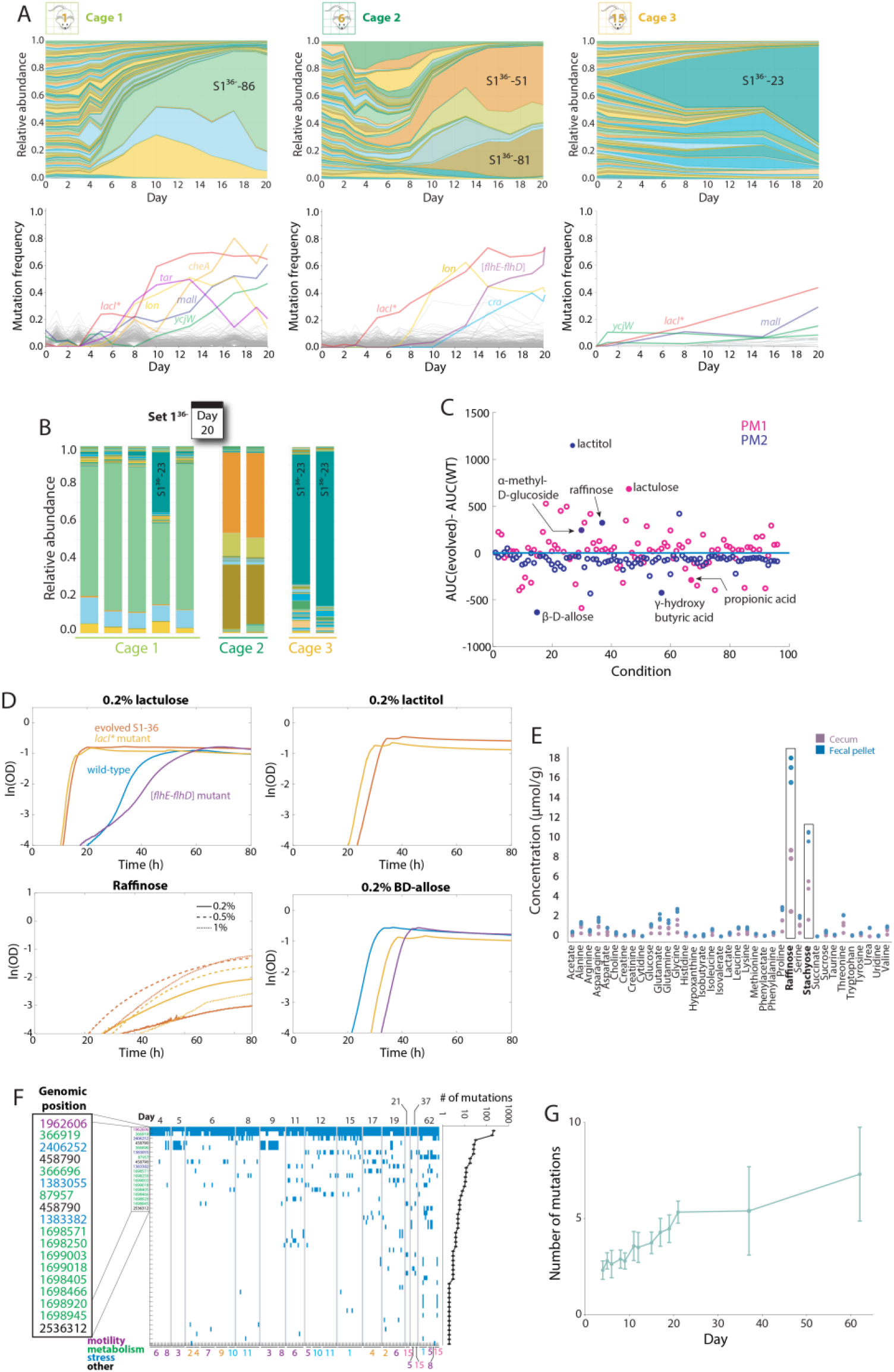
Motility deletions and *lacI* mutations are repeatedly selected for in the mouse gut, in part driven by the presence of raffinose, an abundant carbon source in the gut. A) All S1 strains except S1-36 were used to gavage 9 germ-free mice in three cages on day 0 and tracked via daily fecal sampling for 37 days. Plots of the relative abundances (top) and mutations identified from metagenomic sequencing (bottom) of the barcoded *E. coli* populations from a representative mouse in each cage over the first 20 days of colonization. Different barcodes emerged in each cage, with similar mutations in cage 1 and 2 but not in cage 3. B) By day 20, relative abundances of the S1^36-^ barcodes were similar within each cage, despite the differences across cages. C) The parental wild-type and the evolved S1-36 isolate were inoculated into Biolog plates containing M9 medium with one carbon source per well and grown for 48 h in a plate reader. Shown are the differences between the area under the curve for growth curves of the evolved S1-36 and the parent in each well. Filled circles highlight carbon sources that exhibited the largest differences and/or displayed distinct growth dynamics between the two strains. D) Growth curves of the wild-type parent, evolved S1-36, a *lacI*\* mutant, and an *[flhD-flhE*] mutant in M9 supplemented with four carbon sources. In lactulose, lactitol, and raffinose, evolved S1-36 and the *lacI*\* mutant grew much faster and with a shorter lag than the parent or the [*flhD-flhE*] mutant; in lactitol and raffinose, the parent and [*flhD-flhE*] mutant could not grow at all, suggesting that the ability to utilize raffinose is conferred to the evolved S1-36 by the *lacI*\* mutation. In β-D allose, the parent grew better than the *lacI* or [*flhD-flhE*] mutants; evolved S1-36 exhibited no growth and hence is not shown. E) The concentrations of various amino acids and carbon sources in the feces and ceca of germ-free mice were measured using NMR. Feces and ceca exhibited high concentrations of raffinose and stachyose (*n*=3 mice). F) Map of mutations identified via whole-genome sequencing of 213 S1-36 clones isolated from various mice throughout the experiment. The most prevalent mutations, which include the [*flhE-flhD*] deletion (purple) and mutations mostly in metabolism-related genes (green), are labeled with their genome position. G) The number of mutations detected in S1-36 clones increases over the duration of the experiment, suggesting continued selection. Data are mean values and error bars represent 1 standard deviation, with *n*=3-38 isolates on each day.

After 20 days, the relative abundances of all barcodes were again similar across mice in each cage (Figure 3B). Interestingly, the most abundant strain from cage 3 (S1^36-^-23) also emerged by day 5 in one mouse in cage 1 (#4) (Figure S3B), suggesting transfer of this strain between cages (Figure 3A, right). Notably, the sets of barcodes that expanded in abundance were essentially nonoverlapping with the first experiment or across cages, illustrating the stochastic nature of evolution with regard to the barcoded strains that take over. Mirroring our first experiment, by day 20 the three cages contained alleles at >10% frequency in motility, metabolism, and stress-response genes (Figure 3A, S3B-D), including motility operon deletions of various sizes, and mutations in *cra*, *lon, malI, ycjW*, and *lacI*. The most abundant strain in cage 2 (S1^36-^-51) contained the *lacI*\* and motility deletion [*flhE-flhD*] allele combination identified in our first experiment (Figure 3B). Moreover, the barcoded strain that took over by day 20 in cage 1 (S1^36-^-86) contained the same mutations in *ycjW* observed in the first experiment, as well as *lacI\** (Figure 3B, left and middle). As before, we observed substantial overlap in allele frequencies among mice in the same cage, likely reflecting transmission within cages and takeover of higher-fitness mutants. We also observed some allele overlap between cages, potentially reflecting convergent evolution across cages.

### Metabolic profiling reveals fitness advantages of evolved strains in specific carbon sources

The S1-36 strain dominated in our first experiment (Figure 2A-C), presumably due to a large fitness advantage, and the same alleles emerged in the second experiment (Figure 3A). To identify phenotypes associated with the mutations present in the S1-36 (and other) strains, we introduced the *lacI\** allele into the parent MG1655 strain, and isolated an S1-36 clone with only the [*flhE-flhD*] allele from an early time point. These single-mutation strains were compared phenotypically with the parent and an evolved S1-36 isolate from day 37. To determine if the mutations conferred metabolic advantages, we inoculated the four strains into Biolog phenotype array plates containing minimal media spanning a broad range of carbon sources, and monitored optical density over time. The evolved and *lacI*\* strains grew better than the parent in several conditions, including lactulose, lactitol, α-methyl-D-glucoside, and D-raffinose (Figure 3C). There were also a few conditions in which the parent grew better than the evolved strain, including propionic acid, γ-hydroxybutyric acid, and β-D-allose (Figure 3C). The [*flhE-flhD*] mutant behaved similarly to the parent in all conditions measured. We verified the phenotypes by growing cells in M9 minimal medium with each of the carbon sources as the sole nutrient (Figure 3D).

To ascertain the levels of these carbon sources in the gastrointestinal tract, we collected fecal pellets from germ-free mice fed the same diet as in our monocolonization experiments and then immediately sacrificed the mice and dissected their ceca. We processed the ceca and fecal pellets using NMR to quantify the abundance of several key metabolites (Barroso-Batista et al., 2020). This analysis revealed high concentrations of raffinose and stachyose (Figure 3E), which are non-digestible dietary carbohydrates fermented by many normal gut commensals. Neither the parent nor the three mutants grew on stachyose, indicating that this carbon source did not provide a niche to *E. coli* (Figure S3E). Thus, we conclude that diet-derived raffinose likely provided the growth advantage to the S1-36 strain via the *lacI*\* mutation.

### Metabolic mutations accumulate over time

Since the S1-36 strain began to take over early on (by day 5) in most S1 mice and metagenomic sequencing showed that the *lacI*\* and motility mutations arose approximately concurrently (Figure 2A-C), we sought to characterize genetic changes across the population with greater resolution. We isolated >750 colonies from a range of mice across experimental time points reflecting various notable blooms of specific barcodes, including many in which S1-36 dominated. In total, we ended up with 213 S1-36 isolates across 12 mice and 13 time points. We performed whole-genome sequencing of all isolates to determine allele frequencies and co-occurrence between alleles (Table S2).

All S1-36 isolates contained the [*flhE-flhD*] motility deletion (Figure 3F). Many isolates also carried the *lacI*\* mutation, but 86 clones from three mice instead had a point mutation at a different genomic position in *lacI* (Figure 3F). All but one of these 86 clones also had a short insertion between the *clpX* and *lon* genes, both of which encode proteases (Figure 3F). The two *lacI* mutations were mutually exclusive, and clones from the two classes cohabited some mice (Figure 3F). Moreover, six clones from days 6-12 lacked any *lacI* mutations (Figure 3F), suggesting that the motility deletion was the first to emerge. Of the remaining alleles present in >10 isolates, the vast majority were in the metabolic genes *malI* (8 alleles), *yjcW*, and *cra* (Figure 3F), consistent with metagenomic sequencing (Figure 2A-C). The number of mutations in S1-36 isolates increased steadily as a function of the time point from which they were isolated, with an average of 7 mutations per clone by day 62 (Figure 3G). Thus, we conclude that metabolic genes are a frequent target for mutations.

### Migration during cross-housing rapidly remodels strain abundances and allows quantification of migration rates

The separation of barcodes into two sets afforded the opportunity to study migration of gut bacteria between hosts. In the first experiment, on day 19 we moved two S1 mice and two S2 mice into a new cage and cross-housed them for 18 days (Figure 4A) to quantify the ability of S1 strains to colonize S2 mice and vice versa. After the initiation of cross-housing, both sets of mice experienced invasion by barcoded *E. coli* from the opposite set (Figure 4B). In S1 mice, S1-36 remained dominant, with the total abundance of S2 strains accounting for ~5-10% relative abundance on day 21 and decreasing thereafter (Figure 4B). In S2 mice, S1-36 steadily increased in relative abundance after cross-housing, reaching >80% in all S2 mice by day 28 (Figure 4B, right). These dynamics demonstrate the potential for rapid migration, presumably due to coprophagy. S1-36 increased in abundance at similar rates in the two S2 cross-housed mice (Figure 4B, S4A), suggesting a large fitness advantage relative to all S2 strains.

**Figure 4:**
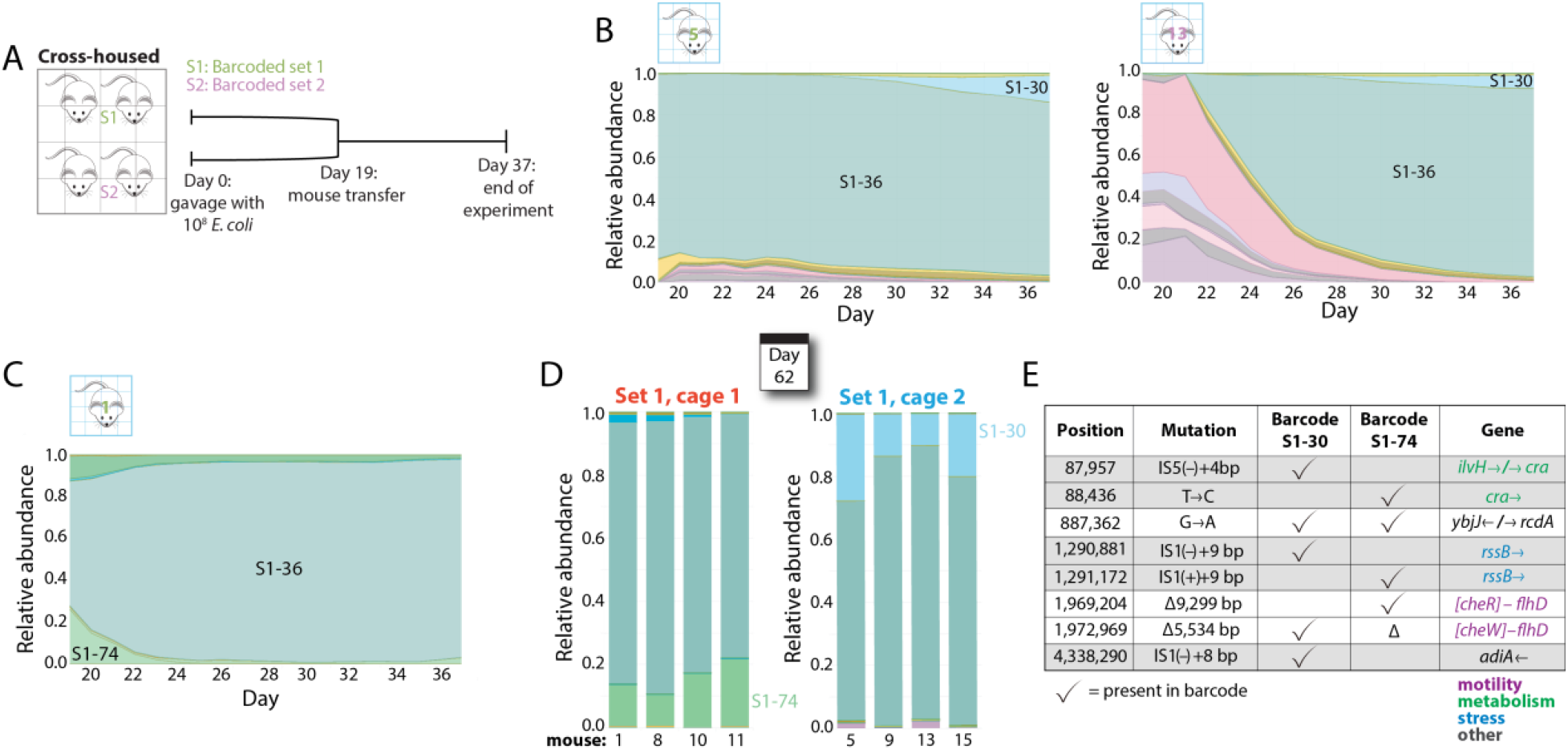
After cross-housing, the dominant strain in S1 mice invades and takes over in all S2 mice. A) Schematic of cross-housing of S1 and S2 mice. Mice colonized with S1 strains were separately housed from mice colonized with S2 strains for 19 days, then transferred into the same cage. Fecal pellets were collected daily until day 37. B) Relative abundances of S1 (shades of green) and S2 (shades of pink) barcodes after day 19 in a mouse initially colonized with S1 strains (left) or S2 strains (right). S1-36 remained dominant in the S1 mouse, and progressively took over the S2 mouse in ~2 weeks. C) In S1 mice that continued to be co-housed with only S1 mice after day 19, strain S1-36 remained dominant on day 37. D) However, by day 62, several other barcodes had started to increase in relative abundance, suggesting continued evolution. E) Strains accompanying strain S1-36 in S1 mice on day 62 contained mutations in many of the same genes, suggesting convergent evolution.

In the S1 mice that remained co-housed with each other after day 19, S1-36 stayed highly abundant (Figure 4C). Unlike the bloom of S1-30 that occurred in the cross-housed mice, S1-74 instead increased in relative abundance, emphasizing the stochasticity of evolution across independent replicates (Figure 4D). On day 62, S1-74 remained the second most abundant strain and had increased to 10-20% in all mice (Figure 4D, left). Similarly, on day 62 the relative abundance of S1-30 in the cross-housed mice was 10-27% (Figure 4D, right). These data suggest that S1-30 and S1-74 developed small fitness advantages over S1-36 that led to slow enrichment over several weeks. Whole-genome sequencing of S1-30 and S1-74 isolates from day 62 revealed that ~80% of the mutations were located in the same genes, while only one mutation was the same between the isolates (Figure 4E), pointing to convergent evolution. Thus, cross-housing effectively displaced the S2 strains, and the S1 strains continued to acquire similar adaptive mutations as the non-cross-housed mice.

### Population-genetics model of fitness advantages in cross-housed mice quantifies the magnitude of migration

During our cross-housing experiment, ~10% of bacteria in S1 mice had S2 barcodes one day after cross-housing (day 20); this fraction subsequently decreased in abundance (Figure 4B). From these data, we inferred that transmission (likely due to coprophagy) typically introduces a sizeable fraction from environmental reservoirs into each host, and that the evolved S1-36 strain had a substantial fitness advantage over the evolved S2 strains. To further interrogate the contributions of transmission and selection to community composition, and to estimate the underlying rates of migration and selection, we developed a population-genetics model to fit the observed barcode abundance dynamics within and across mice that share the same cage. Motivated by our observations above, we assumed that these metapopulation dynamics could be described by a generalized ‘island model’ (Latter, 1973):

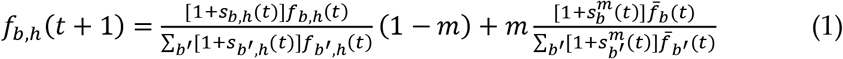

which captures the interplay between local competition within hosts and migration from common environmental reservoir (Figure 5A). In this model, *f_b,h_*(*t*) represents the frequency of barcode *b* in mouse *h* at day *t*, 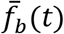 is the average frequency across all mice, and *m* is the overall migration rate; the *s_b,h_*(*t*) and 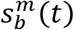 terms represent the fitness differences between barcodes during within-host competition and migration, respectively. These latter quantities will generally vary over time (and across mice) as mutations accumulate within the different barcoded lineages. If all barcodes have the same fitness, this model predicts that the frequencies of barcodes within individual mice will equilibrate within a cage on a timescale of ~1/ln(1 – *m*) days, which increases sharply as *m* approaches zero (Figure 5B). When barcodes differ in fitness, this equilibration must compete with the local amplification of barcodes via natural selection, which occurs over a timescale ~1/s. Thus, the emergent evolutionary dynamics within the metapopulation will strongly depend on the relative magnitudes of *m* and *s_b,h_*(*t*).

**Figure 5:**
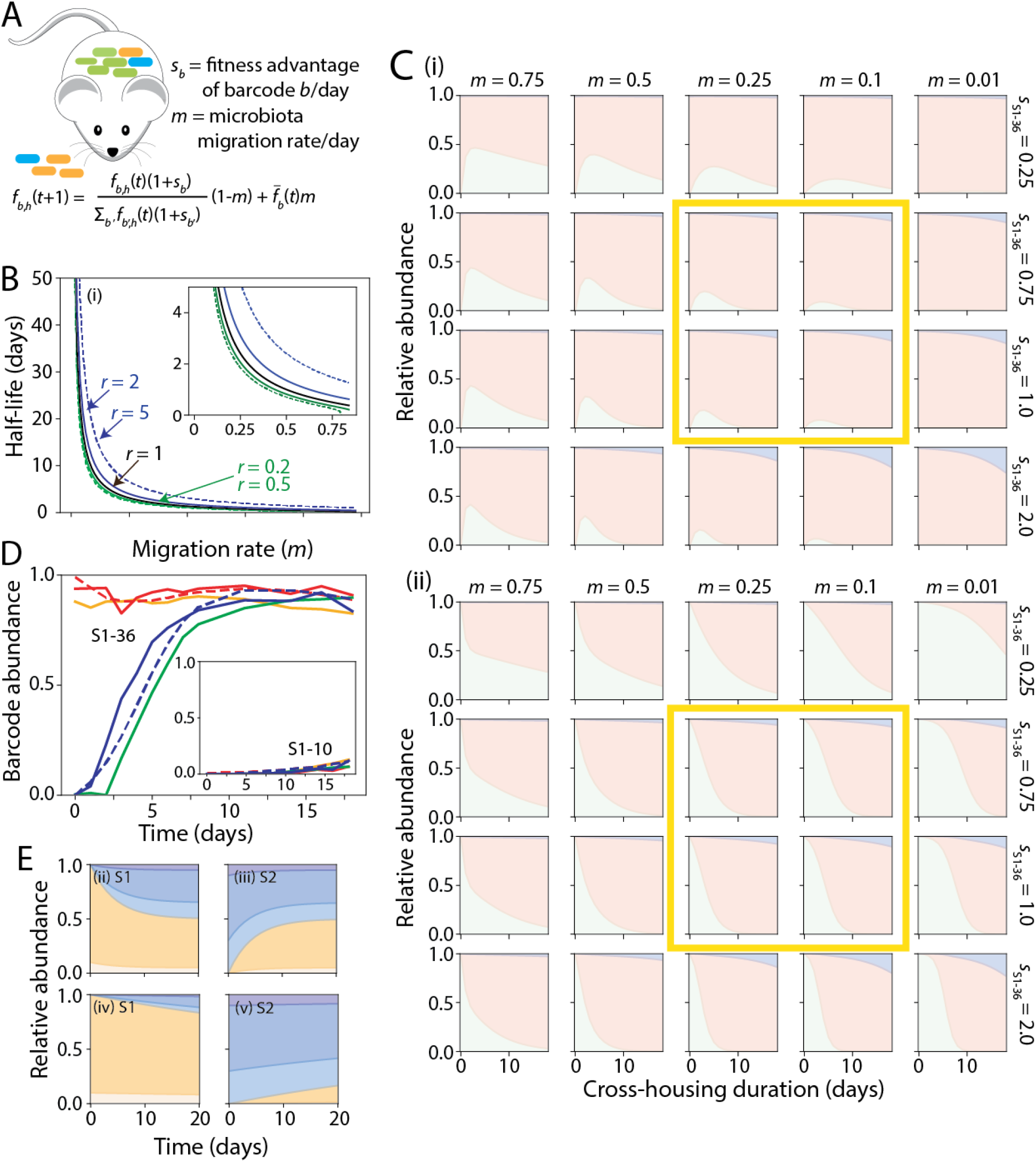
Population-genetics model recapitulates experimental cross-housing data and predicts a high rate of migration each day. A) In the model, the frequency of barcode *b* in mouse *h* on day *t*+1 *(fb,h(t+1))* is determined by the fitness (excess growth/day) of barcode *b* compared with other barcodes (*sb*) and the microbiota replacement fraction each day due to coprophagy 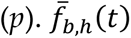 is the relative abundance of barcode *b* on day *t* averaged over all mice, which is assumed to represent the relative load of barcode *b* in feces that is taken in via coprophagy. B) When mice harboring distinct barcode abundances are cross-housed, the model predicts that coprophagy drives all mice to the same barcode composition on a timescale (half-life) that increases strongly with the environmental microbiota replacement rate *m*. In the absence of fitness advantages, the time scale is ~1/ln(1 – *m*) (black line). If mice are predisposed to consume their own feces *r* times as often as the feces of other mice, then the half-life is not strongly affected if 5>*r*>0.2 (green and blue lines). C) Simulations of the model with three barcodes, one each representing S1-36 (pink) and S1-10 (blue) and one representing all S2 barcodes (green), recapitulated experimental data (Figure 3A,B) when *f*_S1-36_ = 0.75 – 1.0, *f*_S1-10_ = 1.33 *f*_S1-36′_, and *m* = 0.1 – 0.25 (yellow boxes). Inset shows the predicted and actual abundances of S1-10 from the time of cross-housing. D) Parameter estimates *f*_S1-36_ = 0.75, *f*_S1-10_ = 1.0, and *m* = 0.12 were obtained from the simulations in (C) by minimizing the squared deviation of the model (dashed lines) from the dynamics of barcodes S1-36 and S1-10 averaged across the four mice in Fig. 3A,B (S1: solid red and orange lines; S2: solid blue and green lines). E) Simulations show that the equilibrium composition is reached within 20 days for two mice with highly distinct starting compositions (i, ii) when *m* = 0.2 and with no fitness advantages among barcodes, but not when *m* = 0.02 (iii,iv).

Estimating these parameters can be challenging due to the time-dependent nature of the within-host fitnesses, *s_b,h_*(*t*), which can potentially vary among hosts in an idiosyncratic way due to clonal interference and frequency-dependent selection. Fortunately, our cross-housing experiment provides an opportunity to directly measure the migration rate, independently of *s_b, h_*(*t*), by focusing on the invasion of new barcodes immediately after cross-housing. For example, on the first day after cross-housing (*t_c_*), the frequencies of S2 barcodes in S1 mice can be directly attributed to migration:

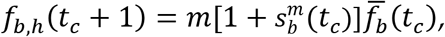

which allows us to infer the relevant migration rates through a simple application of linear regression. We fit this model to our cross-housing data, in which essentially all S2 barcodes migrated into the S1 mice (Figure S5A). Typical migration rates were on the order of 10% per day across a broad range of initial frequencies, suggesting that stochastic transmission bottlenecks do not play a major role in this frequency range (*f*>0.001). These data also revealed some global variation in *m* across mice, as well as some systematic variation in transmission efficiency between barcodes 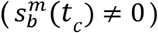. However, higher frequency barcodes did not exhibit systematically higher migration fitness, and the differences between mice averaged out within a day or two (Figure S5B). Together, these data suggest that between-host migration via coprophagy is reasonably well-approximated by an island model with an average migration rate of *m*~10% per day.

Interestingly, these short-term estimates of the migration rate are also qualitatively consistent with observed equilibration timescales in our experiments (Figure 2E, 3B), which typically occurred within ~5-10 days. To make this comparison more explicit, we asked whether we could quantitatively recapitulate our observed data using a coarse-grained version of Eq. 1, in which we assign a constant fitness advantage to a few key lineages. For example, in the cross-housing experiment, we assume that all S2 barcodes and all S1 barcodes except S1-36 and S1-30 had similar (low) fitness and thus could be treated as a single barcode. The fits to barcode abundances in both S1 and S2 cross-housed mice were excellent for *s*_s1-36_ = 0.75,*s*_s1-30_ = 1.0, suggesting that the S1-36 and S1-30 genotypes confer a ~75% and ~100% growth advantage per day, respectively, and *m* = 0.12 (Figure 5C,D), a similar estimate for migration rate in the first few days after cross-housing (Figure S5) and as obtained from simulating the equilibration of S2 co-housed mice (Figure 5E). Predictions based on these parameters (Figure 5C) remained consistent with barcode abundances at day 62 (Figure 4D), suggesting that any subsequent mutations did not dramatically affect the relative fitness advantage of S1-30 over S1-36. Taken together, our dynamic barcode measurements combined with mathematical modeling permitted robust quantitative estimates of key parameters dictating assembly and evolution within mouse intestines.

### Evolution continues after recolonization with a strain of increased fitness

Since strain S1-36 became dominant so quickly in our first experiment, we were hampered in our ability to use the barcode sequences to track further evolution among the population. To overcome this obstacle, we isolated an S1-36 clone on day 37 and cured it of its plasmid, and transformed the resulting strain with 86 S2 plasmids. This new set of strains, which we refer to as S2* strains, allowed us to restart the evolutionary process from a higher-fitness genotype (Table S4). We gavaged three cages of germ-free mice (*n*=10) with the S2* strains and tracked them using barcode and metagenomic sequencing for 43 days. Distinct barcodes dominated the three cages, yet mutations in the same genes emerged, all related to metabolism or motility (Figure 6A). After 20 days, the dominant strain in cage 1 (S2*-62) had mutations in *ycjW*, *malI*, and an intergenic region near *lrhA* (Figure 6A, left), all of which also occurred in S1^36-^ mice (Figure 3A). Interestingly, in mouse 5, strain S2*-69 carried a distinct mutation in *malI* and was at high abundance after day 10; this strain was not displaced by S2*-62 but did not transfer into any of the other mice in the cage (Figure 6B, cage 1). Thus, some alleles may confer fitness in a particular niche that precludes effective transmission.

**Figure 6:**
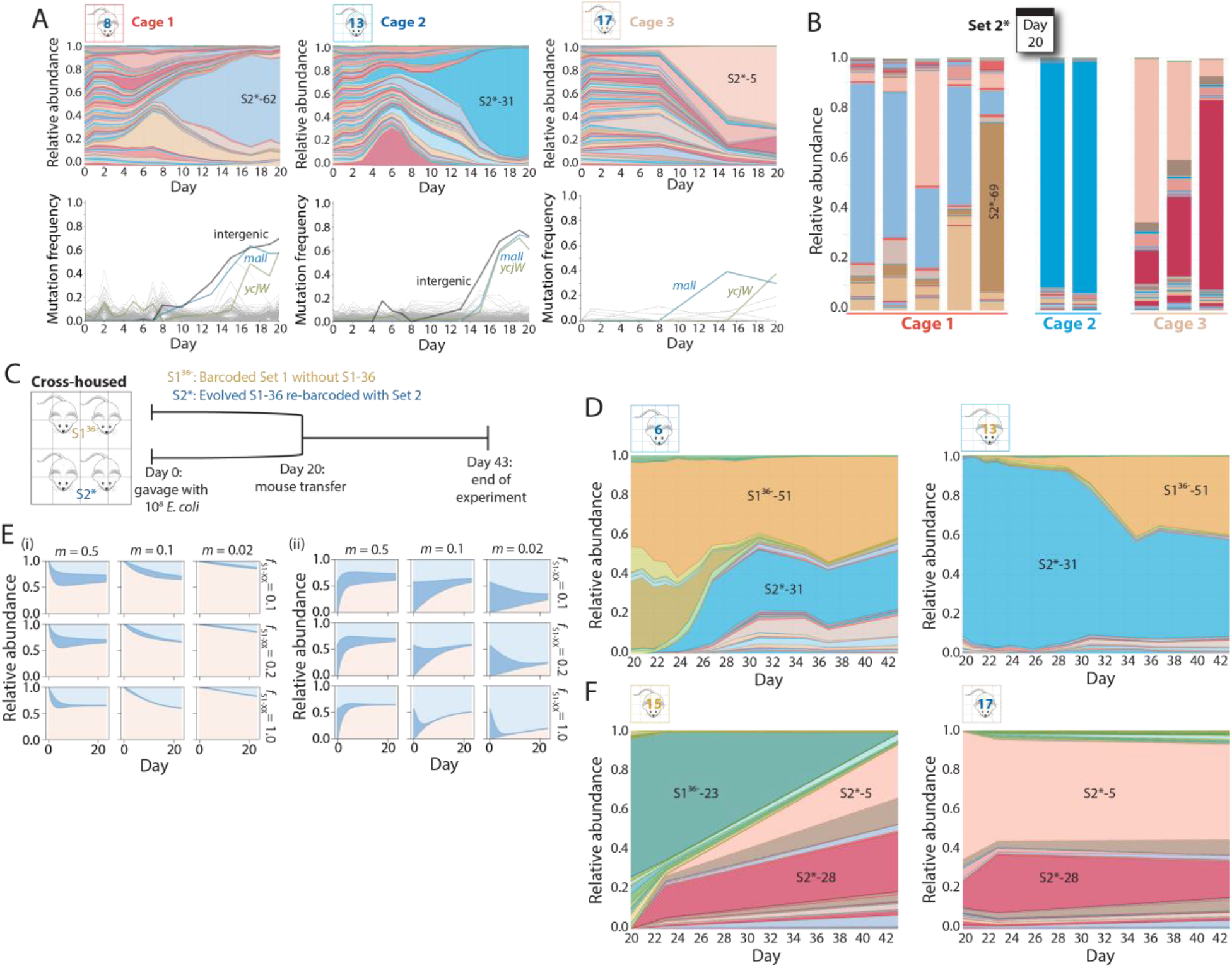
Cross-housing of mice repeatably demonstrates the strength of the motility and *lacI* allele combination. A) S2 barcodes were reintroduced into an evolved clone of S1-36 (S2* strains) used to gavage 10 germ-free mice in three cages on day 0, and tracked via daily fecal sampling for 37 days. Plots of the relative abundances (top) and mutations identified from metagenomic sequencing (bottom) of the barcoded *E. coli* populations from a representative mouse in each cage over the first 20 days of colonization. Various barcodes emerged in each cage, but all three cages hosted bacteria with similar mutations. B) By day 20, the relative abundances of the S2* barcodes were similar within each cage, despite the differences across cages. C) Schematic of cross-housing of S1^36-^ and S2* mice from day 20 to day 43. D) Relative abundances of all barcodes in representative cross-housed S1^36-^ (left) and S2* (right) mice from cage 1. In the S1^36-^ mouse, S2* barcodes invaded and stabilized at ~50% relative abundance, and vice versa for the S2* mouse, suggesting that the two barcode sets have approximately equal fitness. E) The population-genetics model applied to four cross-housed mice with three barcodes, one representing S1^36-^-51 (light blue), one representing all S2* barcodes (pink) with the same fitness as S1^36-^-51, and one representing all other S1 barcodes (blue), recapitulates the coexistence of S1^36-^ and S2* barcodes (D) when *s*_S1-51_ = *s*_s2_ = 0.1 – 1.0 and *m* ≈ 0.1 – 0.25, illustrating the general applicability of the model. F) Relative abundances of all barcodes in representative cross-housed S1^36-^ (left) and S2* (right) mice from cage 2. In the S1^36-^ mouse, S2* barcodes invaded and took over by day 43, while the S2* barcodes were able to maintain colonization in the S2* mice, indicating S2* barcodes were more fit than the S1^36-^ barcodes in this cage.

Our initial cross-housing experiment with S1 and S2 mice combined strains that had evolved for similar amounts of time, but had nevertheless acquired different sets of mutations. By contrast, the high degree of overlap in mutations present in bacteria within mice S1^36-^ and S2* mice led us to hypothesize that their cross-housing could lead to distinct outcomes compared to the cross-housing of the S1 and S2 mice above; note that the S2* colonization experiments were carried out at the same time as the S1^36-^ colonization experiments. To test this hypothesis, we performed two independent cross-housing experiments with S1^36-^ and S2* mice from different cages starting on day 20. We tracked barcode dynamics and metagenomics for an additional 17 days (Figure 6C). These data revealed dramatically different outcomes in the two experiments.

In the first case, strain S1^36-^-51 was at ~50% relative abundance at the time of cross-housing in mice 6 and 7, and remained at ~50% or slightly higher throughout cross-housing (Figure 6D, left; S6B). Meanwhile, the S2* barcodes increased to ~50% in the S1^36-^ mice (Figure 6D, left; S6B); conversely, in cross-housed S2* mice, strain S1^36-^-51 increased over the first ~10 days and stabilized at ~50% (Figure 6D, right). These dynamics suggested coexistence of strains from the two sets with similar fitness, and indeed metagenomic sequencing indicated that S1^36-^-51 had the *lacI*\* mutation, along with *[cheA-flhD]* and mutations in *malI* and *ycjW*, similar to the S2* strains. Thus, we assumed that S1^36-^-51 and all persistent S2* barcodes had the same fitness advantage over all other S1^36-^ barcodes, and used our model to simulate the dynamics of three strains representing S1^36-^-51, all other S1^36-^ strains, and all high-fitness S2* strains. The data were consistent with *s*_s1-51_ ≈ *s_S2_* ≈ 0.2, (Figure 6E), indicating that the motility operon deletions in other S2* strains narrowed the fitness difference compared with the parental strain, against which S1^36-^-51 had a fitness benefit >0.75. While frequency-dependent selection could also play a role in the expansion leading up to coexistence, it is notable that these data were best fit with a similar value for migration rate (*m*~0.1) as in our other co-housing and cross-housing experiments.

The second experiment with cross-housing of S1^36-^ and S2* mice led to a qualitatively distinct outcome from the coexistence in the first experiment. In this case, the S2* strains displayed a clear fitness advantage over the S1^36-^ strains (Figure 6F, S6C), which contained the *lacI\** mutation but not a large motility deletion (Figure 3A, mouse 15). These data suggest that the *lacI*\* mutation alone is not able to compete with strains that also have the [*flhE-flhD*] allele. Taken together, these data illustrate the importance of particular allele combinations in determining coexistence or out-competition.

### Treatment with ciprofloxacin decreases intrastrain diversity and stimulates transmission

Antibiotic treatment causes a massive disruption to the mammalian microbiota that can vary across bacterial species (Ng et al., 2019). To understand how such a disruption would affect bacterial population dynamics at the intra-species level, we treated five S1 mice and two S2 mice with 5 mg of ciprofloxacin once daily on days 19, 20, and 21 (Figure 7A). This treatment effectively reduces the number of bacteria in humanized mice (Ng et al., 2019). To probe the effects of an antibiotic perturbation on transmission, on the same day as the first ciprofloxacin treatment, two S1 mice and two S2 mice were moved into a new cage and cross-housed for 18 days (Figure 7A, bottom). These two cages shared a different isolator from non-antibiotic-treated mice.

**Figure 7:**
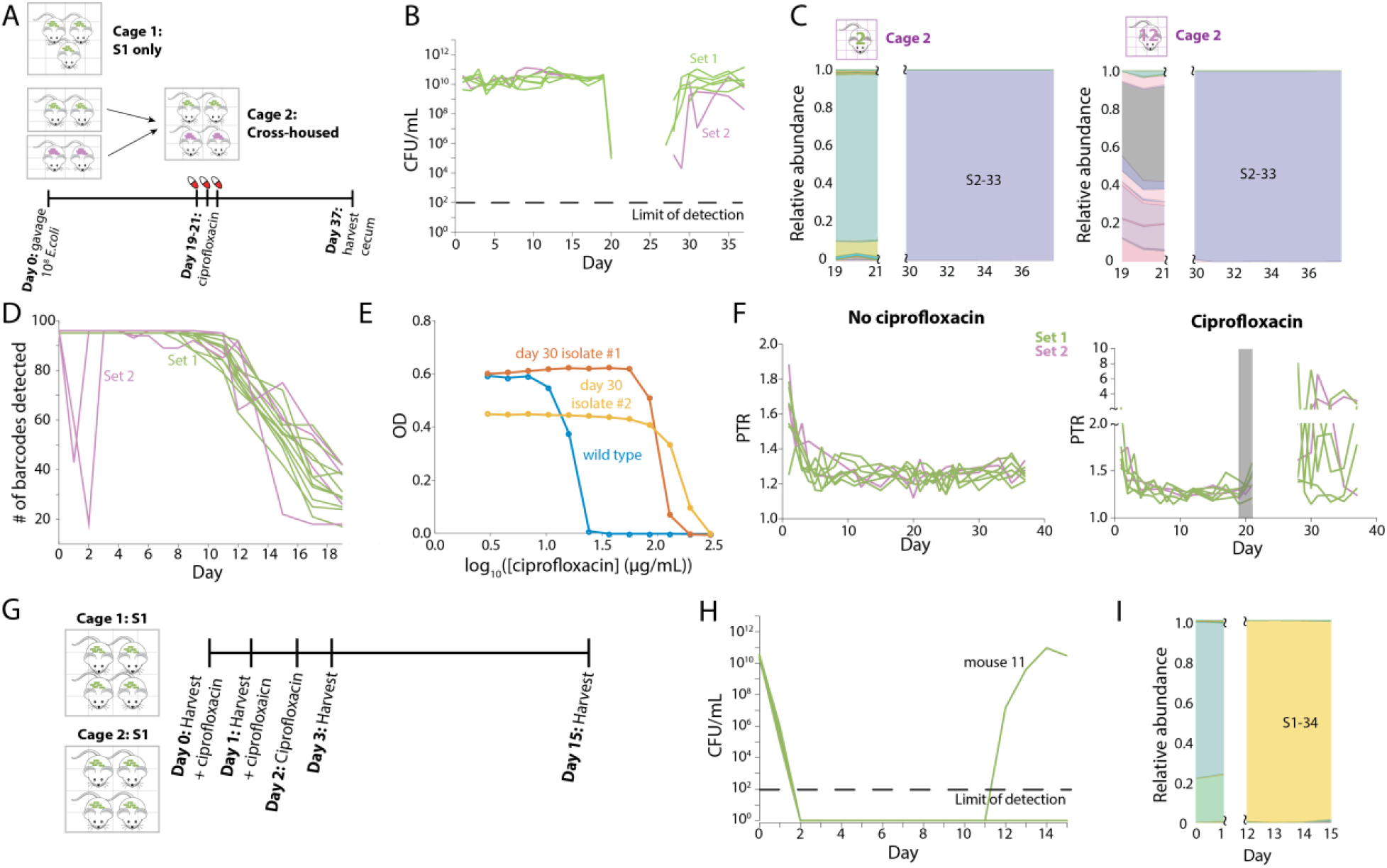
Ciprofloxacin treatment leads to complete loss of substrain-level genetic variation. A) Schematic of ciprofloxacin treatment protocol involving five and two germ-free mice gavaged with S1 and S2 strains, respectively, on day 0. On day 19, two S1 mice were cross-housed with two S2 mice in cage 2. At the same time, the mice in cage 1 (the remaining three S1 mice) and cage 2 began three days of ciprofloxacin treatment. B) CFUs/mL decreased below the limit of detection during ciprofloxacin treatment and did not recover until several days after treatment ended. C) S2-33 was the only strain that recovered in both cages of mice. Omitted days represent days in which bacterial load was too low for accurate sequencing (and hence results resembled water controls). D) S2 mice with uneven barcode abundance distributions on day 1 (Figure S2D) exhibited a large decrease in the total number of detectable barcodes on days 1 and 2, signifying a bottleneck. These two mice quickly recovered the other barcodes, presumably due to coprophagy. All other mice maintained the full set of barcodes until day ~5, when barcodes such as S1-36 started to become dominant. E) Two S2-33 clones isolated on day 30 from mouse 14 displayed higher IC_50_ values than wild type, indicating decreased susceptibility. One clone also grew to lower maximum OD_600_, suggesting a growth tradeoff for the resistance mutation. F) Metagenomic data were used to calculate the PTR in ciprofloxacin-treated (left) and untreated (right) mice over the 37 days of the experiment. Gray rectangle represents the time period of ciprofloxacin treatment. Data for the 7 days after treatment are not shown due to low read counts (Figure S7A). G) Schematic of second ciprofloxacin treatment and harvest protocol in which 8 mice were colonized with S1 barcodes for 2 months before ciprofloxacin treatment daily on days 0, 1, and 2. Mice were also harvested on days 0, 1, and 2, leaving two singly housed mice for tracking until day 15. H) Only mouse 11 showed bacterial recovery, starting on day 12. The other mouse was likely returned to a germ-free state. I) S1-34 was the only strain that recovered in mouse 11.

Culturable densities decreased sharply to ~10^4^-10^6^ CFUs/mL after one day of ciprofloxacin treatment (Figure 7B), and then dropped below the limit of detection (10^2^ CFUs/mL) and remained undetectable for ~7 days in all mice. Barcode sequencing coverage also decreased during this time window, coincident with the drop in density (Figure S7A). Around day 28 (7 days after the last dose), *E. coli* densities began to recover, and by the end of the experiment on day 37 most mice had recovered to pre-antibiotic levels of ~10^10^-10^11^ CFUs/mL (Figure 7B). As soon as densities recovered, the population in all cross-housed mice was dominated by strain S2-33 (Figure 7C, left). Interestingly, strain S2-33 also took over after antibiotic treatment in the cage of only S1 mice (Figure 7C, right), indicating the potential for contamination between cages during antibiotic treatment, presumably due to the opening of spatial or metabolic niches when the resident bacterial population is small.

Sequencing of the S2-33 clones on day 37 revealed a resistance mutation in *gyrA* (D87G), which encodes a subunit of the type II in topoisomerase DNA gyrase. To our surprise, we found that S2-33 clones isolated from as early as day 1 contained this same mutation, suggesting that this resistant strain had persisted at low frequencies in the S2 mice since the beginning of the experiment. We also found that another barcode (S2-90) contained a missense mutation in *gyrB* (E466D) on day 1. No other mutations were identified in these clones (Table S2). Relative to the parent, the IC_50_ to ciprofloxacin increased by 5- and 3.5-fold in the S2-33 and S2-90 isolate, respectively (Figure S7B), indicating a decrease in sensitivity. Sequencing of the initial S2-33 and S2-90 stocks did not identify any mutations in *gyrA* or *gyrB* (or the rest of the genome).

As noted above, the initial barcode unevenness in mice 13 and 15 (Figure S2D) suggested the occurrence of a bottleneck during colonization. Indeed, barcode sequencing revealed a rapid decrease in the number of detectable barcodes and a concomitant increase in the variability of S2 barcode abundance. On day 1, a few barcodes (including S2-33 and S2-90) accounted for most of the abundance in these mice (Figure S2D), and 53 and 37 barcodes were undetectable (<0.01%) (Figure S2D). The observation that distinct gyrase-related mutations arose specifically in these two strains suggested that the conditions that generated the strong bottleneck also selected for gyrase mutants. To test whether the bottleneck was due to acid stresses in the stomach experienced during initial colonization, we grew S2-33 and S2-90 strains isolated from days 1 and 4, respectively, in LB and then resuspended cells in minimal medium without a carbon source at a range of pH values <7. Colony counting indicated that viability after 9 h was the same as the parent across the entire pH range (Figure S7C). Since mutations in topoisomerases have previously been associated with adaptation to high osmolarity (Higgins et al., 1988), we grew the gyrase mutants in LB supplemented with a range of concentrations of the osmolyte sucrose. Again, we observed no advantage (Figure S7D); in unsupplemented LB, the lag time of the two mutants during aerobic growth was actually longer than that of wild type (Figure S7E). Thus, the origin of the bottleneck in these two mice remains unknown. Nonetheless, by day 3, all 96 barcodes were detectable in both of the mice that experienced a bottleneck (Figure 7D), suggesting that environmental reservoirs re-supplied the other strains. The number of barcodes in all S2 mice thereafter followed the same trajectory (Figure 7D). Thus, bottlenecks (likely created by host physiological differences) can select for gyrase mutations that confer increased resistance to ciprofloxacin in the absence of drug, but microbiota composition quickly returns to the expected trajectory in the absence of bottlenecks.

Given the pre-existence of a mutation that decreased ciprofloxacin sensitivity, the prolonged recovery period was presumably due to the retention of high levels of ciprofloxacin in the feces after treatment (Ng et al., 2019), which was likely extended due to the absence of a microbiota that would decrease the antibiotic concentration. We isolated S2-33 clones from mice on day 30 and found that their IC_50_ was several fold higher than the parent (Figure 7E), comparable to the original *gyrA^D87G^* mutant (Figure S7B). Interestingly, IC_50_ values differed somewhat between two S2-33 isolates from mouse 14, and the maximum absorbance was substantially lower in the isolate with the higher IC_50_ (Figure 7E); whole genome sequencing revealed that the mutant with higher IC_50_ possessed additional mutations in *soxR* and *ygiM*. Thus, ciprofloxacin treatment selected for a single barcode, based on its pre-existing resistance to this antibiotic, but with multiple genotypes.

Recent studies have utilized the ratio of the number of metagenomic reads at the origin compared with the terminus as a proxy for the rate of replication, with fast-growing cells having a higher ‘peak-to-trough ratio’ (PTR) (Brown et al., 2016; Korem et al., 2015). Using our metagenomics data, we calculated the PTR for all mice over time. PTR decreased gradually from ~1.8 on day 1 (corresponding to a growth rate of ~1 h^-1^ for *in vitro* culturing (Korem et al., 2015)) to ~1.2 by day 5 (corresponding to early stationary phase *in vitro* (Korem et al., 2015)) (Figure 7F), consistent with the rapid initial increase in CFUs/mL. After ciprofloxacin treatment, concurrent with the recovery in CFUs, the PTR jumped to >2 in most mice, in some cases remaining high for several days (Figure 7F). Thus, recovery after treatment likely signifies higher growth rates during re-expansion. These data demonstrate the wide range of *E. coli* growth rates *in vivo* depending on whether the microbiota is at steady state or recovering from a perturbation.

In the ciprofloxacin experiments above, all mice were in an isolator with S2 mice (even if they were in a different cage) and hence were exposed to S2-33 (which harbored pre-existing gyrase mutations) prior to treatment. Thus, we repeated ciprofloxacin treatment in the absence of S2 strains using the eight S1 mice from the initial experiment that were tracked long-term without ciprofloxacin treatment. On day 62 (when the mice had 9-13 barcodes remaining), we began treatment with 5 mg ciprofloxacin daily for 3 days (Figure 7G). Given the dramatic reduction in CFUs in our first experiment, we sought to determine whether bacteria were still present in the gut and/or cecum but were not being excreted in the feces. We sacrificed six of the eight mice on days 0, 1, 3 during ciprofloxacin treatment, and harvested and stained the large intestine with a FISH probe for Eubacteria. Staining did not detect any bacteria in the large intestine, and we were unable to grow any colonies from the large intestine or cecum (data not shown).

For the remaining two mice (which were singly housed after day 3), we tracked CFUs/mL in the feces daily. In one mouse, no colonies were detected for 13 days after ciprofloxacin treatment (Figure 7G). To test whether there were reservoirs of viable *E. coli* within the gut that did not get incorporated into feces, we sacrificed the mouse on day 15 and ground up the entire intestine before plating. We did not detect any colonies (data not shown), indicating that these mice were likely returned to a germ-free-like state by ciprofloxacin treatment.

In the other mouse, the recovery time was substantially longer than our previous experiment (Figure 7B), with CFUs/mL only increasing above 10^2^ ten days after treatment ended (day 12) and stabilizing at 10^10^-10^11^ thereafter (Figure 7H). The population consisted of a single barcode (S1-34) after day 12 (Figure 7I), and this strain showed no prior sign of fitness advantage: levels were below the limit of detection (<0.01%) on days prior to antibiotic treatment (Figure 7I). Whole-genome sequencing revealed a *gyrA^A119K^* mutation that was distinct from those of the S2 barcodes from the first experiment and conferred an even larger increase in IC_50_ (Figure S7F). The mutation was likely already present at very low levels at the onset of treatment, as it would have been >10,000 times more likely for a *de novo* mutation to emerge in the more abundant barcodes instead. The extended interval before recovery may be due to the lower population sizes of S1-34, or to the absence of coprophagic exchange with other hosts due to single housing that would reinforce recovery. Regardless, these data highlight the possibility for recovery even when poised at the boundary of extinction.

## Discussion

Here, we have showcased the power of DNA barcodes for tracking the evolution of nearly 200 *E. coli* strains during the colonization of a live mammalian host. These strains were introduced in initially equal numbers, and selection distorted the distribution within days (Figure 1). The plasmids were maintained in the absence of stabilizing antibiotic for at least seven days (Figure 2), indicating that other, antibiotic-sensitive bacteria can be introduced to track barcode dynamics within a multispecies community. In the future, introducing a larger number of barcoded bacterial strains into mice may permit the quantification of the fitness of more barcoded strains (Levy et al., 2015) prior to population takeover. Nonetheless, the rapid dynamics that we observed in this study suggest that a large fraction of the barcodes may be driven extinct within a few weeks regardless of the initial library size.

Barcodes can be employed to study processes that are otherwise difficult to analyze due to the inability to distinguish phenotypically and/or genotypically identical populations. Here, we used barcoding to quantify the extent and effects of microbial transmission between mammalian hosts. The fortuitous finding that certain S2 mice had dramatically different initial barcode distributions (Figure S2D) positioned us to determine that transmission of bacteria between mice in the same cage (presumably via coprophagy) caused barcode abundances to become highly similar within ~1 week.

Using a population-genetics model to fit barcode dynamics of co-housed mice and after cross-housing of mice colonized with different libraries (Figure 5), we inferred the fitness benefits of key mutations and the magnitude of migration, through which ~10-20% of the resident *E. coli* population is introduced exogenously each day. Importantly, the model was sensitive to parameter values, and consistently predicted similar values across all of our experimental data (Figure 5, 6E). These findings suggest that coprophagy is a strong homogenizing force on the microbiota, such that the microbiota of mice in the same cage are highly non-independent. It is likely that migration rates are lower in humans, given the lack of coprophagy; this study suggests that singly housed mice may be better than co-housed mice for modeling the human microbiota. Future studies of barcoding in other commensals, particularly obligate anaerobes, will be useful to reveal whether the same degree of homogenization occurs when survival in the environment (due to the presence of oxygen) is more challenging than it is for the facultative anaerobe *E. coli*. In the future, barcoding could also be used to shed light on microbial exchange with environmental reservoirs through spike-ins, and to determine whether mice engage in selective coprophagy (e.g. as a function of mouse genotype).

Our metagenomic analyses of >1,300 samples yielded complementary insights into evolutionary dynamics, providing mechanistic insights into why certain barcoded strains took over the community. In some cases, the same mutation emerged in multiple barcoded strains that continued to coexist (Figure 6D), suggesting either that the two strains are spatially segregated but subject to the same selective pressures or the existence of a microecology involving the two strains. In addition, we sometimes observed competing lineages within the same barcode (clonal interference, Figure 2B), presumably due to the rapid and continued evolution of strains throughout our experiments. Disentangling these intra-barcode dynamics was greatly assisted by sequencing of large numbers of isolates, a strategy that has also been exploited during *in vitro* studies (Li et al., 2019). In the two S2 mice that were surprisingly transiently dominated by strains with gyrase mutations (Figure S2D), it is unclear why these gyrase mutations were selected for during colonization (which occurred in the absence of antibiotics), since they do not confer any benefits during normal growth or growth/survival at higher osmolarity or lower pH (Figure S7C,D). Given that all other mice experienced approximately uniform colonization by all barcodes (Figure 1D), it seems likely that the physiology of the two mice with uneven barcode abundances applied a bottleneck with distinct and stronger selective pressures than the other mice during colonization. Regardless, these fortuitous findings indicate that antibiotic resistance can be selected for due to pleiotropic fitness benefits in other environmental conditions.

By coupling daily barcode tracking data and metagenomic sequencing, we discovered rapid and repeatable waves of selection of mutations involved in motility and metabolism (Figure 2A-D, 3A). Remarkably, the selection of large deletions in the flagellar operon coupled to mutations in carbon utilization genes was reproducible in independent mouse experiments (Figure 3A), presumably reflecting the strong selective advantage of this combination of mutations. A previous study showed that deletion of motility genes provides *E. coli* with a growth advantage in the gut by redirecting energy from motility processes to growth and by allowing the expression of genes normally repressed by the *flhDC* operon (Gauger et al., 2007). S2 strains were outcompeted by S1 strains upon cross-housing (Figure 4B), even though the S2 strains carried motility-related and metabolic mutations (Figure 2D); the higher fitness of the *lacI* [flhD-flhE]* genotype in the S1-36 strain likely reflects the greater benefits of the 4-bp *lacI*\* insertion over the *lacI* point mutations in S2 strains.

Our NMR analysis of gut and cecal contents of germ-free mice on a standard diet revealed an abundance of the trisaccharide raffinose (Figure 3E), which the evolved S1-36 strain but not the parental strain had the ability to metabolize. These findings are similar to a recent study demonstrating adaptations that increase *E. coli’s* ability to compete for amino acids during monocolonization (Barroso-Batista et al., 2020). Together, these studies show that nutritional competition is a major selective pressure in intra-species interactions driving *E. coli* evolution in the mouse gut, and suggest (perhaps unsurprisingly) that diet is a strong selective force and that identification of available metabolic niches may be a good predictor of evolutionary dynamics; this idea is consistent with previous findings that porphyran was sufficient to bias the microbiota toward a *Bacteroides* species with exclusive access (Shepherd et al., 2018). The *lacI*\* mutation was selected for in a previous long-term evolution experiment *in vitro* involving passaging in minimal medium with glucose/lactose (Quan et al., 2012); curiously, our measurements indicate that *lacI*\* confers higher fitness on lactitol and lactulose (Figure 3D). Previous efforts successfully selected for raffinose utilization through UV mutagenesis, and determined that expression of the alpha-galactosidase necessary for raffinose utilization requires constitutive beta-galactosidase expression (Lester and Bonner, 1957). We were unable to spontaneously evolve raffinose utilization *in vitro* after a week of passaging, either on plates or in liquid culture, perhaps due to the rare occurrence of the 4-bp insertion. The repeated emergence of the *lacI*\* mutation *in vivo* may be due to larger population sizes *in vivo* than *in vitro*, a larger selective advantage, fewer tradeoffs, and/or a higher mutation rate *in vivo* compared with *in vitro*. Regardless, our findings indicate that the presence of an unoccupied niche such as raffinose utilization can serve as a strong evolutionary driver, motivating future studies of barcoded library colonization of mice (or humans) fed various diets to identify nutrient-dependent changes to the trajectory and rate of evolution.

The high sensitivity of *E. coli* to ciprofloxacin meant that treatment eliminated almost all cells (Figure 7B,H), leading to the selection of a single barcode after a surprisingly long recovery time (Figure 7C,I). These findings indicate that even if certain species appear to recover from antibiotics, intrastrain-level heterogeneity that is important for host and microbiota health may be eliminated during treatment, highlighting yet another negative consequence of antibiotics. Notably, when mice were singly housed, the recovery period was extended relative to co-housed mice (Figure 7H), with one mouse seemingly reverting to a germ-free state. This delay in recovery may be due to the lack of microbiota reinforcement from other mice in the cage; these findings are reminiscent of our recent work showing that single housing leads to heterogeneous antibiotic recovery trajectories (Ng et al., 2019; Oliveira et al., 2020), and further demonstrate the importance of the environmental microbial reservoir. In the future, other antibiotics or perturbations applied to barcoded libraries can be used to quantify the magnitude of disruption necessary to cause a decrease in intrastrain diversity via the extinction of barcoded strains.

In this study, we chose a laboratory strain of *E. coli* to accelerate evolution, presuming that MG1655 has evolved away from the host environment due to its longstanding use as a laboratory model organism. Given the rapid pace of *E. coli* evolution *in vivo*, do other commensals exhibit evolutionary dynamics on a similar timeline, or do these “wild” species already carry the most fit adaptations? Notably, colonization of germ-free mice with a *Bacteroides thetaiotaomicron* transposon library revealed metabolic mutations that increase fitness *in vivo* (Goodman et al., 2009), suggesting the existence of evolutionary potential. However, despite the vast array of monocolonization experiments that have been carried out with various gut commensals, there is very little data on the frequency of adaptive mutations in these species. A second aspect that may affect the pace of evolution is the presence of other species; for example, a recent study found that bicolonization of mice with *E. coli* and *Blautia coccoides* altered the gut metabolome and the evolutionary trajectory (Barroso-Batista et al., 2020). Our results suggest that another scenario that would likely alter the evolutionary trajectory of *E. coli* is if another species such as *Bacteroides ovatus* (Gherardini et al., 1985) can fill the raffinose-utilization niche. Nonetheless, we detected continued evolution of the S1-36 strain after it was re-barcoded and re-introduced into the gut (Figure 6A), indicating that the selective landscape was not saturated. Regardless, the prevalence of metabolic mutations suggests that identification of genetic variation within an individual’s microbiota could provide key information for personalized dietary recommendations.

Overall, our study demonstrates that barcoded libraries will be a powerful resource for interrogating microbiota function in the future; importantly, barcodes provide a high-dimensional window into each species’s evolution, without any obvious phenotypic cost.

The continued development of genetic tools for gut commensals (Dodd et al., 2017; Guo et al., 2019; Shiver et al., 2019) should drive the creation of phylogenetically diverse barcoded libraries. Important insight can arise from coupling barcode and metagenomic dynamics to metabolomics to better understand metabolic selective pressures; given the high degree of repeatability in our experiments, the ability to tune the metabolic environment may allow engineering of a particular evolutionary trajectory. More generally, using barcodes to determine selective forces will reveal important insights into adaptation to the gut environment.

## Acknowledgments

The authors thank the Huang and Sonnenburg labs for helpful discussions. The authors acknowledge support from the Allen Discovery Center at Stanford on Systems Modeling of Infection (to K.C.H.), NIH RM1 Award GM135102 (to K.C.H.), and a Friedrich Wilhelm Bessel Award from the Humboldt Foundation (to K.C.H.). J.L.S. and K.C.H. are Chan Zuckerberg Biohub Investigators. This work was also supported in part by the National Science Foundation under Grant PHYS-1066293 and the hospitality of the Aspen Center for Physics.

