## Supplemental Figures and Methods for "Quantifying the interplay between rapid bacterial evolution within the mouse intestine and transmission between hosts"

### Supplemental Figure Legends

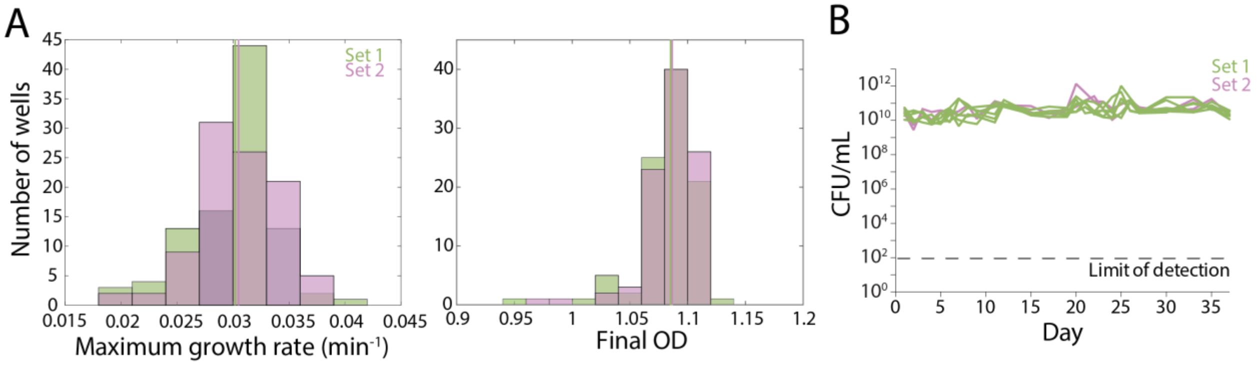


**Figure S1: Growth dynamics of the barcoded *E. coli* libraries, related to Figure 1.**

A) The individual barcoded *E. coli* from Set 1 (green) and Set 2 (pink) showed similar maximum growth rates and final $OD_{600}$ values. Vertical lines represent the mean of each set.

B) Culturable densities of bacteria on LB agar plates from the feces of all mice were approximately constant.

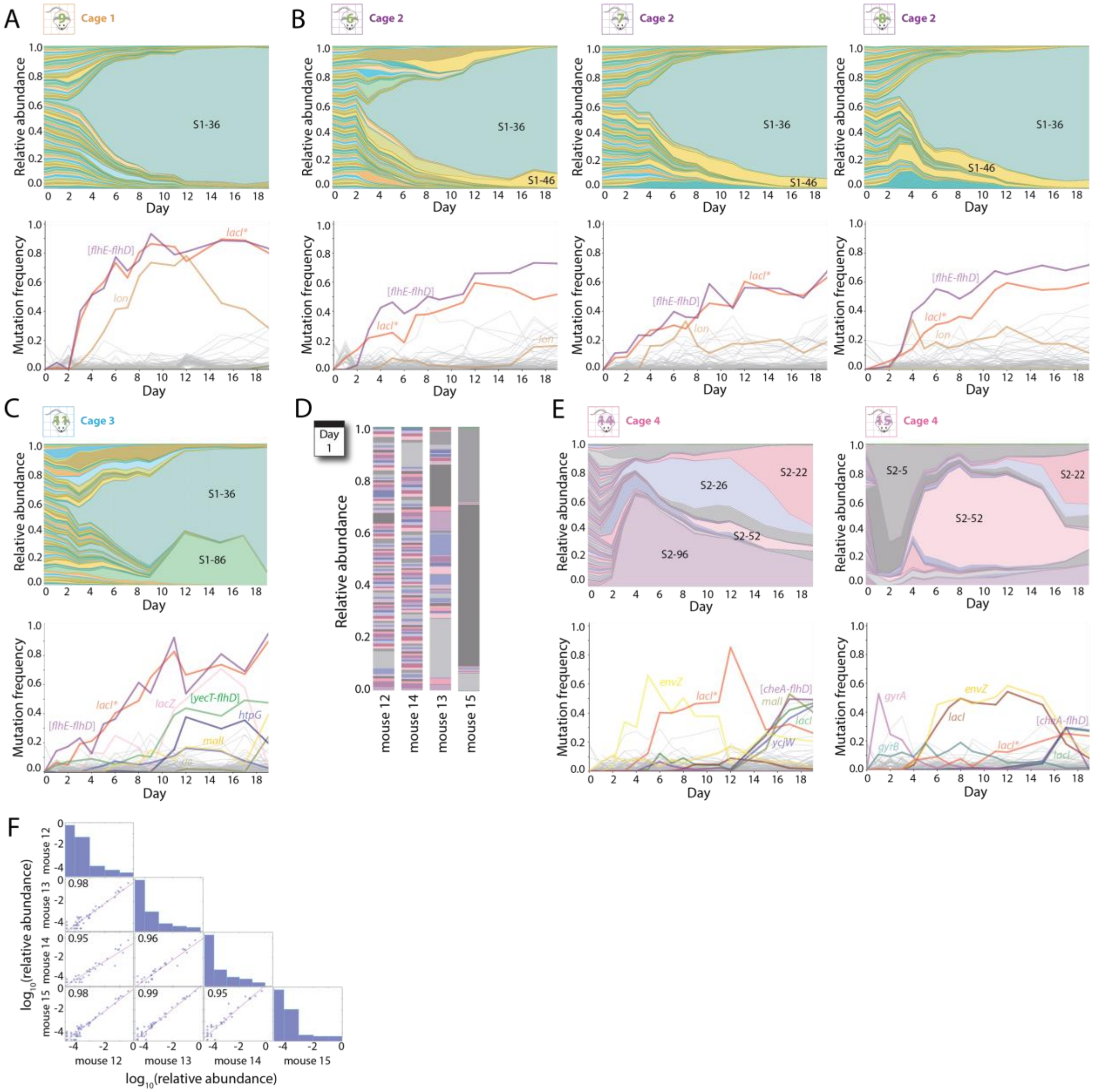


**Figure S2: Additional barcode and metagenomic tracking of S1 and S2 mice, related to Figure 2.**

A-C) Relative abundances of barcodes (top) and mutations (bottom) over the first 19 days in S1 mice not shown in Figure 2.

D) Relative abundances of all S2 mice on day 1 after colonization. An apparent bottleneck was observed in mice 13 and 15 that led to the dominance of a few barcodes, whereas mice 12 and 14 had relatively even abundances.

E) Relative abundances of barcodes (top) and mutations (bottom) over the first 19 days in S2 mice not shown in Figure 2.

F) Pearson correlation coefficients of relative abundances between all pairs of S2 mice were very close to 1 on day 19, indicating similar compositions after 19 days of co-housing. Histograms along the diagonal show the distribution of barcode abundances in each mouse.

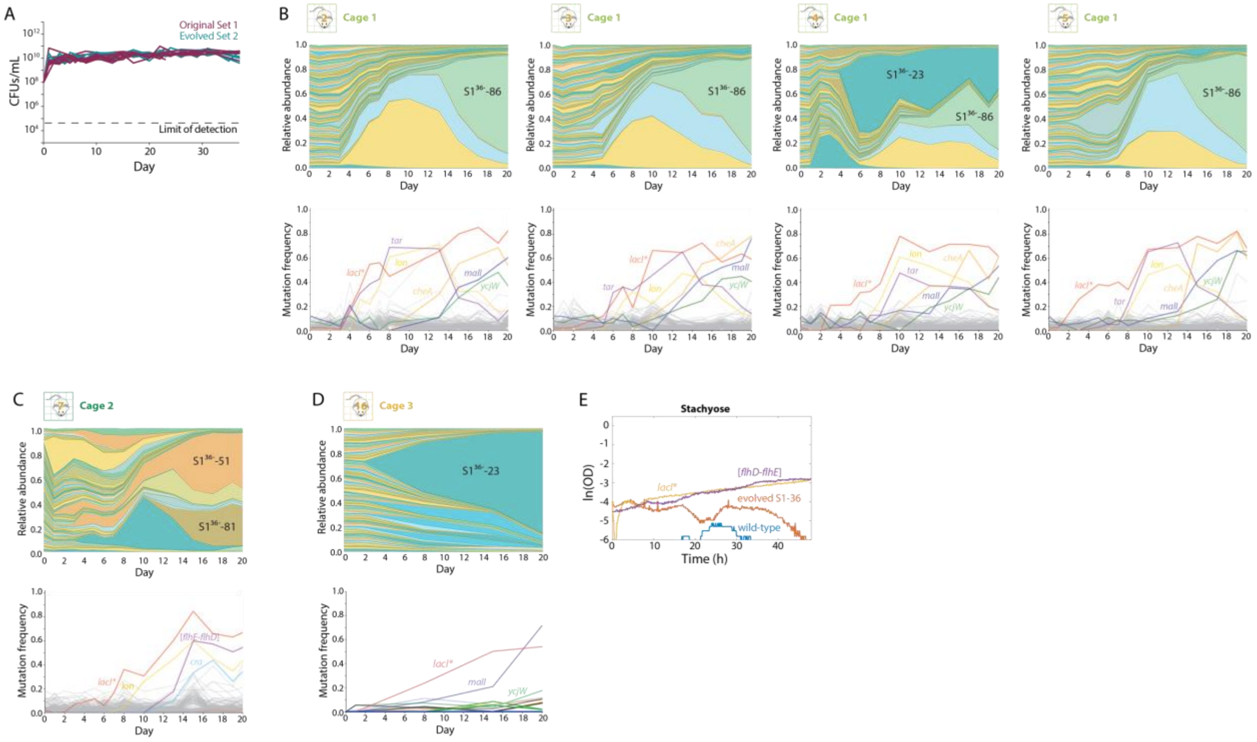


**Figure S3: Additional barcode and metagenomic tracking of $S1^{36-}$ and S2* mice, related to Figure 3.**

A) Culturable densities of bacteria on LB agar plates from the feces of all mice were approximately constant.

B-D) Relative abundances of barcodes (top) and mutations (bottom) over the first 20 days in $S1^{36-}$ mice not shown in Figure 3.

E) Growth curves of the wild-type parent, evolved S1-36, a *lacI** mutant, and an [*flhD-flhE*] mutant in M9 supplemented with stachyose in a Biolog plate.

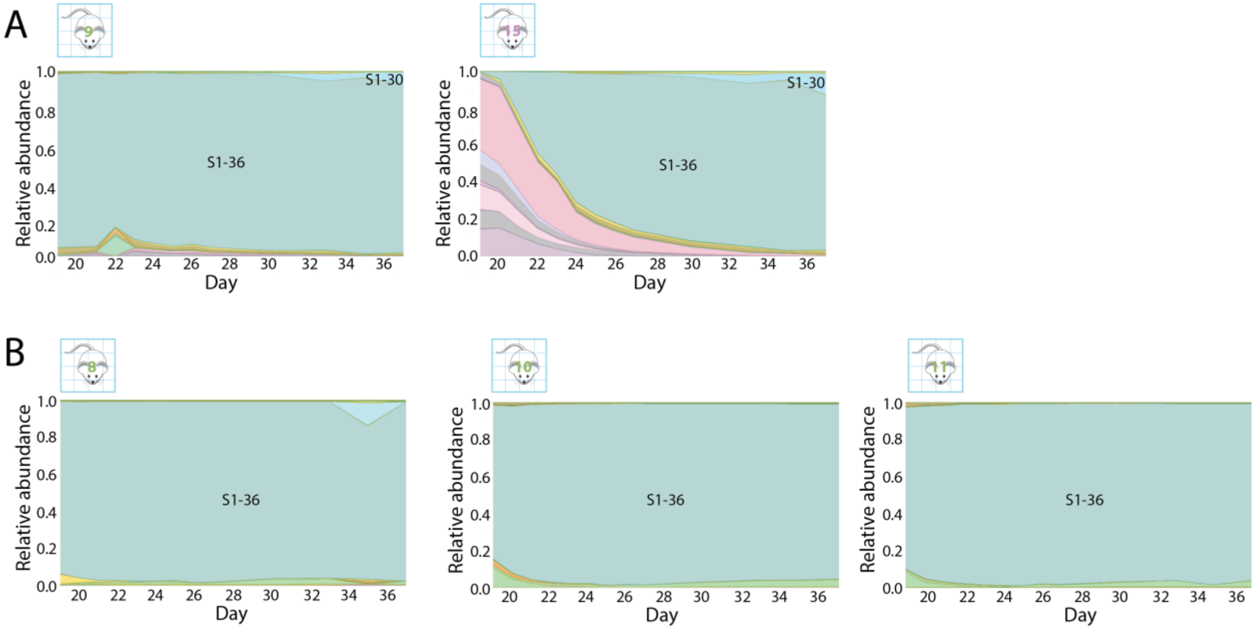


**Figure S4: Additional barcode data from cross-housed S1 and S2 mice, related to Figure 4.**

A) Relative abundances of S1 (shades of green) and S2 (shades of pink) barcodes in after 19 days in mice initially colonized with S1 (left) or S2 strains (right) that are not shown in Figure 4, related to Figure 4B.

B) Relative abundances of S1 barcodes after day 19 in mice initially colonized with S1 strains that were not cross-housed and not shown in Fig. 4.

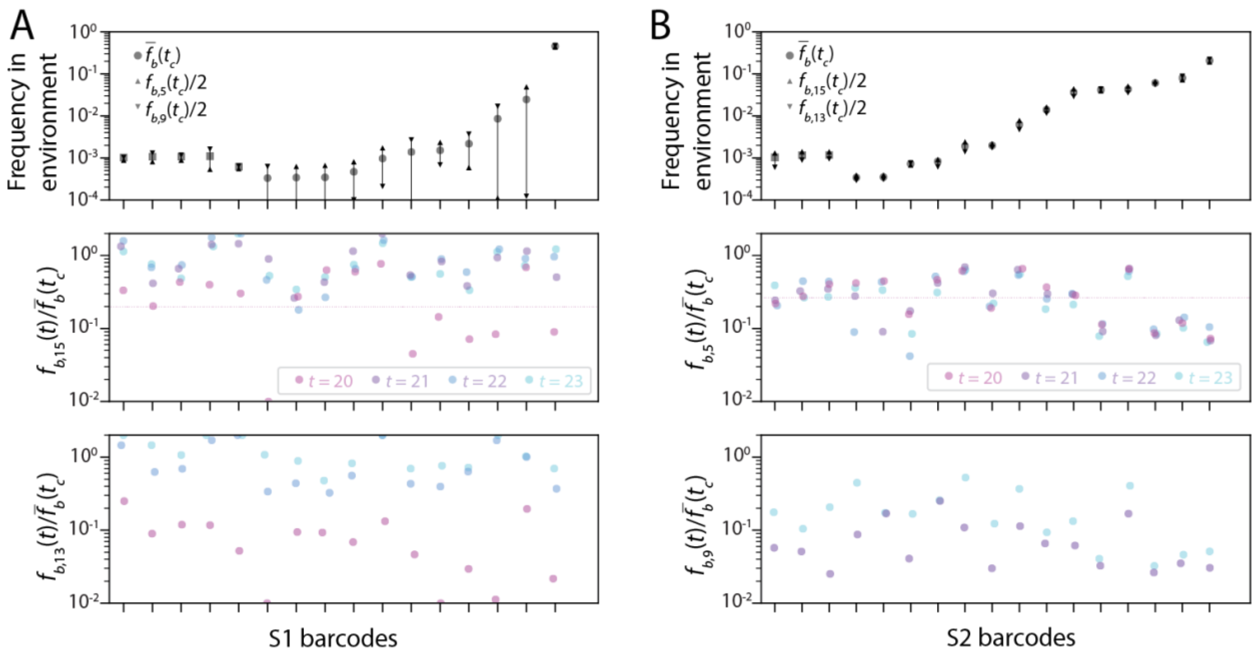


**Figure S5: Cross-housed mouse data supports a migration rate of ~10%, related to Figure 5.**

A) Top: frequencies of the most abundant S1 barcodes present in S1 mice on day 19. Circles represent the mean of individual barcodes across mice 5 and 9, squares represent course-grained barcodes that represent collections of barcodes whose individual frequencies were <$10^{-3}$. Triangles are individual measurements in mice 5 and 9. Middle, bottom: the abundance of the S1 barcodes in S2 mice 15 (middle) and 13 (bottom) on days 20-23 (directly after cross-housing) normalized to the mean abundance in S1 mice at the time of cross-housing ($\bar{f}_b(t_c)$), which represents a time- and barcode-dependent migration rate. Pink dotted line is the mean migration rate on day 20, which is ~10%.

B) Top: frequencies of the most abundant S2 barcodes present in S2 mice on day 19. Circles represent the mean of individual barcodes across mice 15 and 13, squares represent course-grained barcodes that represent collections of barcodes whose individual frequencies were $<10^{-3}$. Triangles are individual measurements in mice 15 and 13. Middle, bottom: the abundance of the S2 barcodes in S1 mice 5 (middle) and 9 (bottom) on days 20-23 (directly after cross-housing) normalized to the mean abundance in S2 mice at the time of cross-housing ($\bar{f}_b(t_c)$). Pink (middle) and purple (bottom) dotted lines are the mean migration rate on day 20 or day 21, respectively, which were ~10% similar to (A).

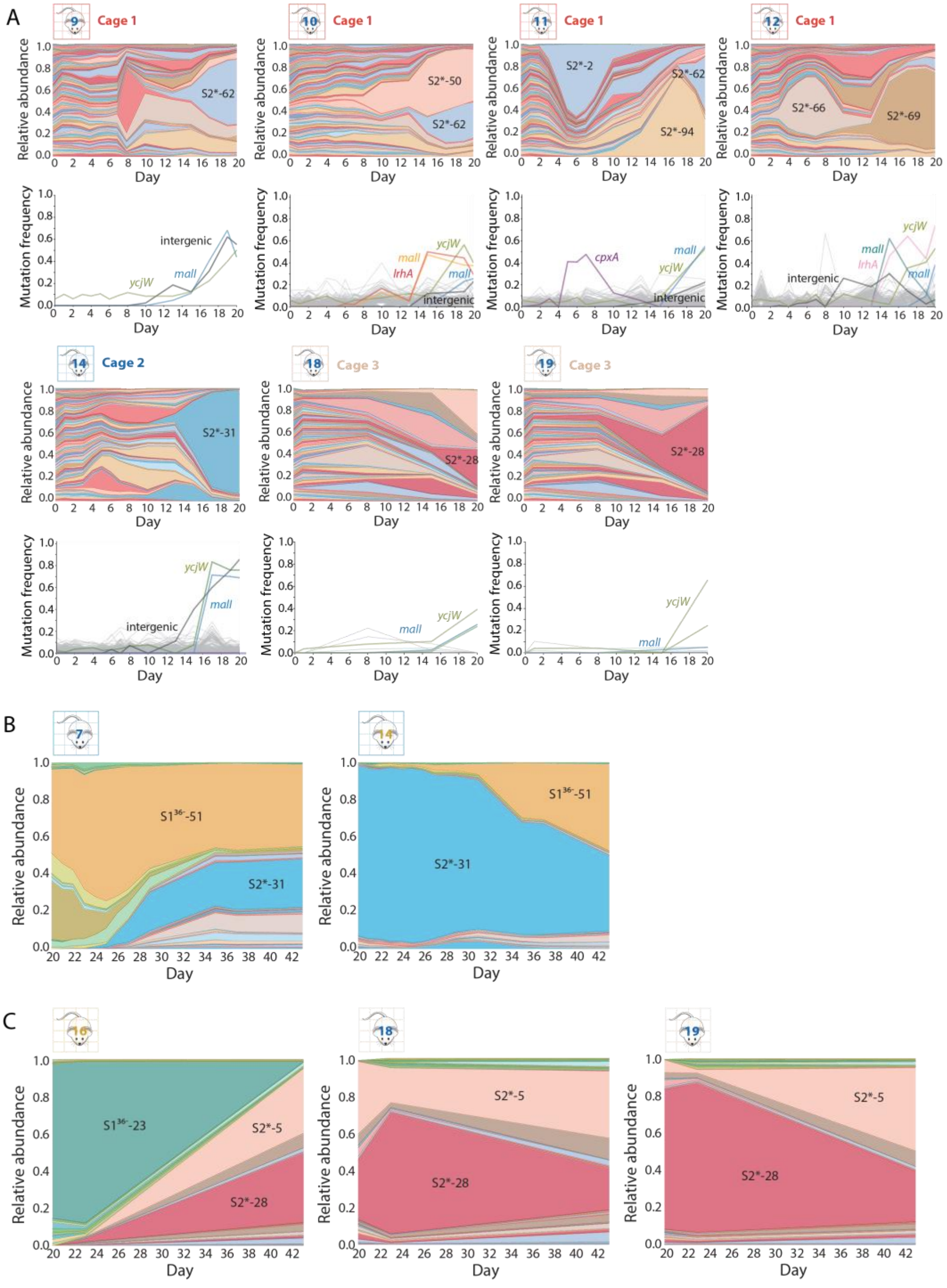


**Figure S6: Additional barcode and metagenomic tracking of $S1^{36-}$ and S2* mice, related to Figure 6.**

A) Relative abundances of barcodes (top) and mutations (bottom) over the first 20 days in S2* mice not shown in Figure 6A.

B) Relative abundances of barcodes after day 20 in cross-housed $S1^{36-}$ and S2* mice not shown in Figure 6D.

C) Relative abundances of barcodes in cross-housed $S1^{36-}$ and S2* mice from cages 2 not shown in Figure 6F.

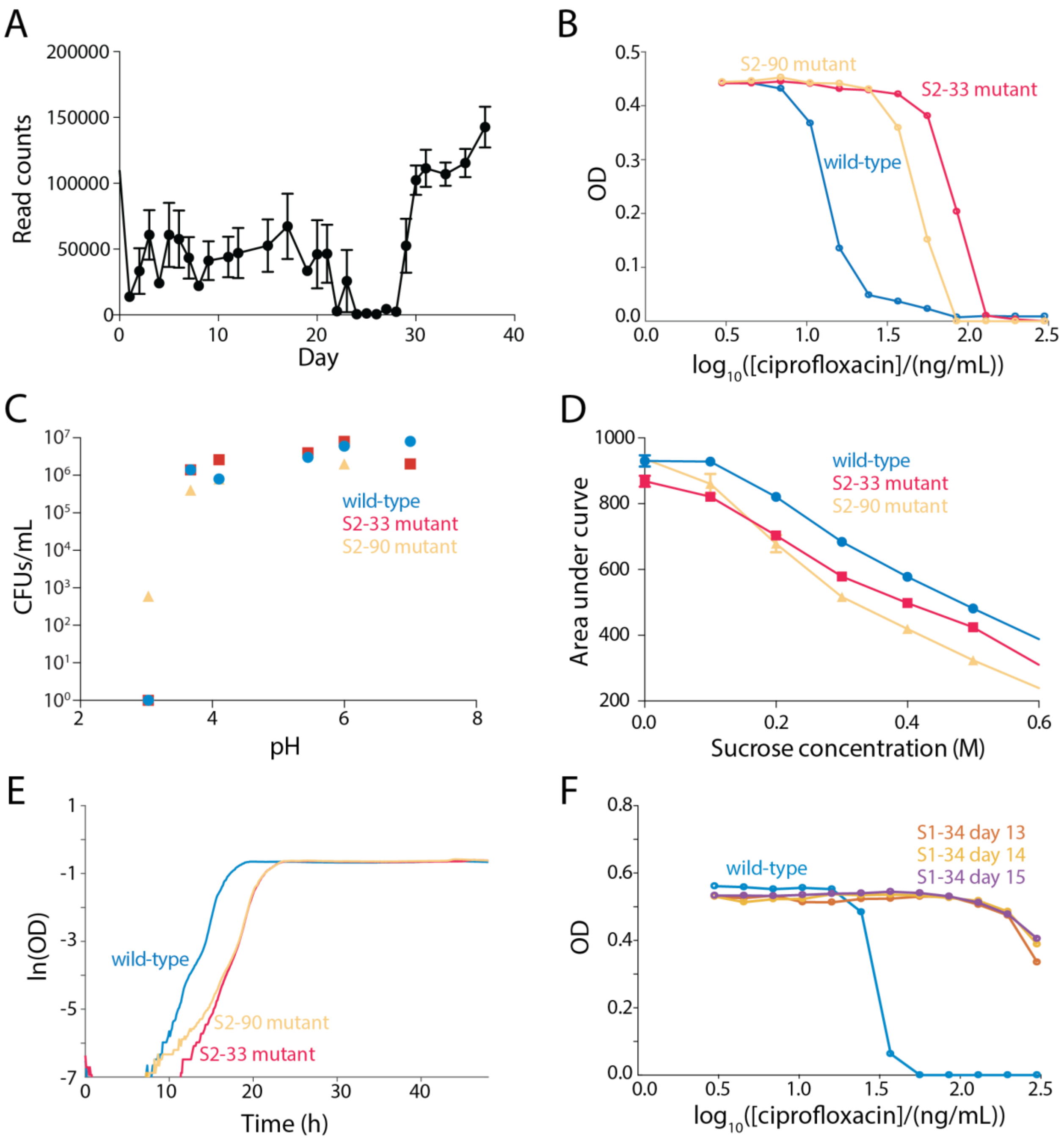


**Figure S7: Growth phenotypes of ciprofloxacin-resistant clones, related to Figure 7.**

A) Read counts from barcode sequencing of all ciprofloxacin-treated mice dropped during treatment (days 19-21) and remained low until day 29. Error bars represent 1 standard error of the mean.

B) The S2-33 *gyrA* mutant and S2-90 *gyrB* mutant exhibited higher $IC_{50}$ to ciprofloxacin than the wild-type parent.

C) Culturable bacterial densities after 24 h in M9 salts at a range of pH values were similar for wild type, the S2-33 *gyrA* mutant, and the S2-90 *gyrB* mutant.

D) Area under the curve after 24 h of growth was slightly higher for wild type compared with the S2-33 *gyrA* and S2-90 *gyrB* mutants across a range of sucrose concentrations.

E) The S2-33 *gyrA* and S2-90 *gyrB* mutants exhibited a longer lag than wild type in unsupplemented LB.

F) Three S1-34 *gyrA* mutants isolated on different days displayed higher $IC_{50}$ than the wildtype parent strain.

### Methods

#### Mouse strains

All mouse experiments were conducted in accordance with the Administrative Panel on Laboratory Animal Care, Stanford University's IACUC. Experiments involved female Swiss-Webster mice 6-12 weeks of age. For all experiments, mice were co-housed unless stated otherwise. For monocolonization experiments, mice were gavaged with $10^8$ cells of an equal mixture of *E. coli* cells and fed normal mouse chow (LabDiet 5010) *ad libitum* throughout the experiment. Mice were euthanized with $CO_2$ and death was confirmed via cervical dislocation.

#### Barcode library creation

Barcodes were generated in (Cira et al., 2018). To generate the barcoded library in *E. coli* MG1655, plasmid DNA was extracted from TOP10 *E. coli* cells (ThermoFisher) with a miniprep kit according to the manufacturer's instructions (Macherey-Nagel). Plasmids were transformed into chemically competent *E. coli* MG1655 cells via heat shock and plated on LB agar plates containing chloramphenicol (35 μg/mL) and ampicillin (100 μg/mL). Colonies were picked, grown separately in LB broth with chloramphenicol (35 μg/mL) and ampicillin (100 μg/mL), and banked as glycerol freezer stocks at -80 °C.

***lacI** mutant creation**

*E. coli* MG1655 cultures were grown from a glycerol stock for 12 h in LB at 30 °C. The next day, cultures were diluted 1:100 and grown at 32 °C to a final $OD_{600}$ of 0.4-0.6. Cultures were incubated at 42 °C for 15 min and then placed in an ice slurry for 5-10 min. Cultures were washed in ice-cold water three times in decreasing volumes of 50 mL, 800 µL, and 200 µL. Fifty microliters of competent cells were mixed with 2 µL of 100 µM oligo (C*C*A* T*TA AGT TCT GTC TCG GCG CGT CTG CGT CTG GCT GGC TGG CTG GCA TAA ATA TCT CAC TCG CAA TCA AAT TCA GCC GAT AGC GGA ACG, where * indicates a phosphorothioated DNA base) and electroporated at 1.8 kV. Finally, cells were rescued in fresh LB for 2 h at 37 °C and plated on LB agar. Mutants were identified via growth on M9+0.2% raffinose and confirmed through Sanger sequencing of the *lacI* gene using primers 5′-TGG CTG GCT GGC TGG CAT AAA T (forward) and 5′-CGC AGC CCG GAC TCG GTA AT (reverse).

**Measurements of population growth**

Cultures were grown from glycerol stock for 12 h in LB with chloramphenicol (35 µg/mL) and ampicillin (100 µg/mL) at 37 °C with constant shaking. Then, cultures were diluted to $5\times10^5$ cells/mL and 200 µL of the dilutions were transferred to clear-bottom transparent 96-well plates. For measurements of antibiotic $IC_{50}$, cells were diluted into fresh LB medium with 3-300 ng/mL ciprofloxacin. Plates were sealed with transparent film pierced with a laser cutter to create ~0.5-mm holes that allowed aeration in each well. Absorbance was measured at 600 nm in an Epoch2 plate reader (BioTek Instruments). Plates were shaken between readings with linear and orbital modes for 145 s each. Growth rates and lag times were quantified using custom code in MATLAB R2016b (Mathworks).

**Plasmid curing of evolved strain**

A gentamicin plasmid with the same origin of replication as the barcoded plasmid was used to transform a culture of *E. coli* S1-36 isolated from a mouse from the first experiment on day 37. Cells were plated on LB agar containing 200 μg/mL gentamicin. Colonies were picked, grown separately in LB broth with 200 μg/mL gentamicin, and replated on LB agar plates containing 35 μg/mL chloramphenicol to determine which colonies had lost the barcoded plasmid. Chloramphenicol-sensitive clones were confirmed to lack the barcoded plasmid through Sanger sequencing. One of these clones was transformed with the set 2 barcoded plasmids as described above, and recovered on LB agar plates containing 35 μg/mL chloramphenicol. 86 out of the 96 barcoded plasmids were successfully used for transformation, creating the S2* strains. Loss of the gentamicin plasmid was achieved by growing the strains in LB with 200 μg/mL gentamicin, and Sanger sequencing confirmed gentamicin plasmid loss.

**DNA extraction and sequencing**

DNA was extracted from whole fecal pellets with PowerSoil and PowerSoil-htp kits (Qiagen). For barcode sequencing, extracted DNA was amplified using a two-stage platinum *Pfx* DNA polymerase PCR (Thermo). In a one-cycle PCR, adapter regions and unique molecular identifiers (barcodes) were added to each template. This step generates one uniquely labeled functional template per initial template plasmid molecule. The initial primers were then removed using Ampure XP Beads at a ratio of 1:1 (Beckman Coulter). A second PCR was run for 35 cycles to amplify these labeled templates. During this reaction, known DNA indices were attached to the product to allow informatic demultiplexing of pooled libraries.

First PCR: 94 °C for 2 min, 2X (94 °C for 15 s, 53 °C for 30 s, 68 °C for 30 s), 68 °C for 5 min, hold at 4 °C.

Primers:

5′-

ACACTCTTTCCCTACACGACGCTCTTCCGATCTNNNNNNNNTCGCTAAGGATG

ATTTCTGGA-3'

5′-

GACTGGAGTTCAGACGTGTGCTCTTCCGATCTNNNNNNNNNNNNNNNNTCGC

TTGGACTCCTGTTGAT-3'

NNN = unique molecular identifier (barcode)

Second PCR: 94 °C for 2 min, 35x (94 °C for 15 s, 72 °C for 30 s, 68 °C for 30 s), 68°C for 5 min, hold at 4 °C.

Primers:

5′-

AATGATACGGCGACCACCGAGATCTACACnnnnnnAAACACTCTTTCCCTACACG

ACGCTCTTCCGATCT-3'

5′-

CAAGCAGAAGACGGCATACGAGATAAnnnnnnGTGACTGGAGTTCAGACGTGTG

CTCTTCCGATCT-3'

nnnnnn = multiplexing indices

Successful reactions were confirmed via agarose gel electrophoresis and pooled in equal abundance. The sequencing library was then finalized via purification with Ampure XP Beads at a ratio of 1:1. Sequencing was performed on an Illumina MiSeq with read length 2×75 bp and an average of 100,000 reads per sample.

For metagenomic and whole-genome sequencing, extracted DNA was arrayed into a 384-well plate and concentrations were quantified and equilibrated using the PicoGreen dsDNA quantitation kit (ThermoFisher). DNA was added to a tagmentation reaction, incubated for 10 min at 55 °C, and immediately neutralized. Mixtures were then added to 10 cycles of a PCR that appended Illumina primers and identification barcodes to allow for mixing of samples during sequencing. Wells were mixed, using 1 μL per well, and the pooled library was purified twice using Ampure XP beads to select the appropriately sized bands. Finally, library concentration was quantified using a Qubit (Thermo Fisher). Sequencing was performed on a NovaSeq S4 with read lengths of 2×146 bp.

**Sequence analysis**

Barcode sequences were analyzed with *SeqPrep* v.1.3.2 (https://github.com/jstjohn/SeqPrep) and custom MATLAB scripts, and plotted using *Tableau Desktop* v. 10.4. Genome sequences were analyzed using a previously described pipeline (Good et al., 2017).

Some barcode sequences that were not included in some samples nevertheless had very small numbers of reads assigned to them, and applying more stringent thresholds by counting only exact matches to species barcodes did not remedy this apparent noise, so it is unlikely that the noise was due to sequencing errors that resulted in misclassification of one species barcode as another. Instead, this noise likely arose from index hopping that occurs when multiplexing samples on Illumina sequencers. Thus, we acknowledged a noise floor below which reads could not be reliably detected. Based on previous work with these barcodes (Cira et al., 2018), we applied a 0.1% threshold for barcode abundances. While this threshold is conservative, it ensures that detected barcodes were actually present in the experiment.

**Quantification of bacterial densities**

Bacterial densities were quantified by spot plating in duplicate on plates with LB only and with LB supplemented with ampicillin (100 μg/mL) and chloramphenicol (35 μg/mL). Plates were incubated in a 37 °C warm room.

**Preparation of mouse fecal pellets for metabolomics**

Cecum and fecal contents of germ-free mice were collected and quickly stored at -80 °C. Samples were processed as reported previously (Barroso-Batista et al., 2020).

**PTR (peak-to-trough ratio) analysis**

Metagenomic sequencing data was used to compute PTR as reported previously (Brown et al., 2016).
